# GAS6-AXL signaling triggers actin remodeling and macropinocytosis that drive cancer cell invasion

**DOI:** 10.1101/2020.03.15.993147

**Authors:** Daria Zdżalik-Bielecka, Agata Poświata, Kamila Kozik, Kamil Jastrzębski, Kay Oliver Schink, Marta Brewińska-Olchowik, Katarzyna Piwocka, Harald Stenmark, Marta Miączyńska

## Abstract

AXL, a member of the TAM (TYRO3, AXL, MER) receptor tyrosine kinase family, and its ligand GAS6 are implicated in oncogenesis and metastasis of many cancer types. However, the exact cellular processes activated by GAS6-AXL remain largely unexplored. Here, we identified an interactome of AXL and revealed its associations with proteins regulating actin dynamics. Consistently, GAS6-mediated AXL activation triggered actin remodeling manifested by peripheral membrane ruffling and circular dorsal ruffles (CDRs). This further promoted macropinocytosis that mediated the internalization of GAS6-AXL complexes and sustained survival of glioblastoma cells grown under glutamine-deprived conditions. GAS6-induced CDRs contributed to focal adhesion (FA) turnover, cell spreading and elongation. Consequently, AXL activation by GAS6 drove invasion of cancer cells in a spheroid model. All these processes required the kinase activity of AXL but not TYRO3, and downstream activation of PI3K. We propose that GAS6-AXL signaling induces multiple actin-driven cytoskeletal rearrangements and macropinocytosis that jointly contribute to cancer cell invasion.

## Introduction

Metastasis, the ability of cancer cells to spread from the primary tumor and invade distant secondary sites, makes cancer incurable. Despite much progress in oncology in the last decades, metastasis still causes approximately 90% of cancer-related deaths (Lambert et al, 2017). To initiate metastasis, cancer cells need first to disassemble cell-cell and cell-substrate adhesion sites and prepare for migration and invasion through the extracellular matrix (ECM), vessels and tissues. This requires, among others, a significant remodeling of the plasma membrane and actin cytoskeleton (Chitty et al, 2018; Lambert et al, 2017).

During migration and invasion, cancer cells form various actin-based protrusions such as lamellipodia, filopodia, invadopodia, membrane blebs and circular dorsal ruffles (CDRs) (Buccione et al, 2004; Caswell & Zech, 2018; Hoon et al, 2012). CDRs are enigmatic actin-rich, ring-shaped structures formed transiently on the dorsal surface of cells, in response to certain growth factors. Up to date, it was demonstrated that CDRs are formed upon stimulation with platelet-derived growth factor (PDGF) in fibroblasts, hepatocyte growth factor (HGF) in HeLa and polarized epithelial MDCK cells, and epidermal growth factor (EGF) in fibroblasts and liver-derived epithelial cells (Abella et al, 2010; Azimifar et al, 2012; Chiasson-MacKenzie et al, 2018; Corallino et al, 2018; Mellstrom et al, 1988). The functions of these structures are still not fully explored, but they have been postulated to play a role in preparation of cells for motility, mesenchymal migration through ECM, and macropinocytosis (Chiasson-MacKenzie et al, 2018).

Macropinocytosis is an evolutionarily conserved, actin-dependent form of endocytosis that mediates non-selective uptake of a large amount of extracellular fluid into cells. During macropinocytosis, peripheral ruffles (PRs) collapse inwards to create large plasma membrane-derived vesicles, termed macropinosomes, which contain extracellular fluid and solutes (Kerr & Teasdale, 2009; Marques et al, 2017). Macropinosomes may also form concomitantly with the contraction and closure of CDRs (Hoon et al, 2012). Generally, macropinocytosis allows rapid and efficient remodeling of the plasma membrane and its composition. Another proposed function of macropinocytosis is to support cellular metabolism, particularly of cancer cells where macropinocytosis is common and can be induced by mutated RAS. It has been shown that pancreatic cancer cells expressing oncogenic RAS upregulate macropinocytosis to acquire extracellular albumin which, upon lysosomal degradation, provides amino acids for the metabolism and cell growth. In this way, macropinocytosis allows cancer cells to survive in a nutrient-poor tumor microenvironment (Commisso et al, 2013; Palm, 2019; Recouvreux & Commisso, 2017).

AXL is a receptor tyrosine kinase (RTK) implicated in oncogenesis. Together with TYRO3 and MER, it belongs to the TAM family. TAM receptors are dispensable for embryonic development but participate in phagocytic clearance of apoptotic cells (efferocytosis) in adult organisms (Lemke, 2013; Lemke & Burstyn-Cohen, 2010; Nguyen et al, 2013). Two known TAM ligands are vitamin K-dependent proteins: growth arrest-specific 6 (GAS6) and anticoagulant protein S (PROS1). GAS6 appears to bind all three TAMs, with the highest affinity for AXL, whereas PROS1 predominantly binds TYRO3 and MER (Lew et al, 2014; Tsou et al, 2014).

AXL is associated with the pathogenesis of a wide array of human cancers, including gliomas, melanomas, breast, lung, gastric, prostate, ovarian and kidney cancer (Paccez et al, 2014; Verma et al, 2011; Wu et al, 2014). Overexpression of AXL and/or GAS6 has been shown to correlate with poorer prognosis and increased cancer invasiveness, for example in glioblastoma patients (Hutterer et al, 2008). Inhibition of AXL signaling reduced glioma cell migration, invasion and proliferation *in vitro* and prolonged survival of mice after intracerebral implantation of glioma cells (Onken et al, 2016; Vajkoczy et al, 2006). An increased expression of AXL in highly metastatic breast cancer was found to be essential for all steps of the metastatic process, starting with intravasation of cancer cells (Goyette et al, 2018). Consistently with the association of AXL with cancer invasion and metastasis, a very recent study of Revach et al (2019) reported that AXL may regulate the formation of invadopodia in melanoma cells. Furthermore, AXL has been linked to epithelial-to-mesenchymal transition (EMT) that is associated with metastasis (Chaffer et al, 2016; Diepenbruck & Christofori, 2016). For example, expression of GAS6 and AXL was upregulated in cells that underwent TGFβ1-induced EMT (Antony & Huang, 2017; Antony et al, 2016; Wilson et al, 2014). Several studies showed that AXL activation, associated with an EMT-like phenotype, conferred resistance to both conventional and targeted anti-cancer therapies (Paccez et al, 2014; Wu et al, 2014). Thus, AXL inhibition constitutes a promising therapeutic strategy (Paccez et al, 2014). Accordingly, R428, a first-in-class AXL kinase inhibitor, is being tested in the second phase of clinical trials for metastatic lung and triple-negative breast cancer, glioblastoma and acute myeloid leukemia (Holt et al, 2018; Schuster et al, 2018; Sheridan, 2013).

Despite multiple roles of AXL in cancer invasion and metastasis, the molecular mechanisms underlying its action in cancer cells are not fully characterized. Here, we identified an interactome of AXL using proximity-dependent biotin identification (BioID) assay. Our results reveal intracellular processes induced by GAS6-AXL signaling and mechanisms underlying GAS6-AXL-driven cell invasion.

## Results

### AXL interacts with proteins implicated in the regulation of actin cytoskeleton

To determine the AXL interactome we performed the BioID assay under basal and ligand-stimulated conditions in HEK293 cells and human glioblastoma LN229 cells, which do not express and express AXL, respectively (Fig 1A). We selected the HEK293 line as it is widely used for the BioID assay (Kim et al, 2014; Lambert et al, 2015; Nishimura et al, 2015; Roux et al, 2012), whereas LN229 cells represent a cancer-relevant model to study GAS6-AXL signaling. To perform proximity labeling, we fused HA-tagged, mutated biotin ligase (BirA-R118G-HA denoted BirA*-HA) to the intracellular C-terminal part of AXL (AXL-BirA*-HA). Expression of fusion proteins BirA*-HA and AXL-BirA*-HA was verified in HEK293 cells (Fig EV1A). Immunofluorescence analysis showed that AXL-BirA*-HA fusion protein localized properly to the plasma membrane (Fig EV1B). AXL-BirA*-HA displayed also activation upon ligand stimulation, as visualized by phosphorylation of the receptor following GAS6 treatment (Fig EV1C). Since these results confirmed the functionality of the AXL-BirA*-HA fusion, we generated HEK293 and LN229 cell lines stably expressing BirA*-HA or AXL-BirA*-HA (Fig 1B, Fig EV1D). These lines were incubated in the presence or absence of GAS6, and with biotin for 24 h, as this incubation time ensured efficient target biotinylation (Fig EV1E). Biotinylated proteins isolated using streptavidin-coated magnetic beads were analyzed by mass spectrometry. Western blot analysis of samples before (input) and after (output) streptavidin-mediated pull-down showed significant enrichment of biotinylated proteins in the output samples, in comparison to the input (Fig 1C, Fig EV1F).

**Fig 1.**
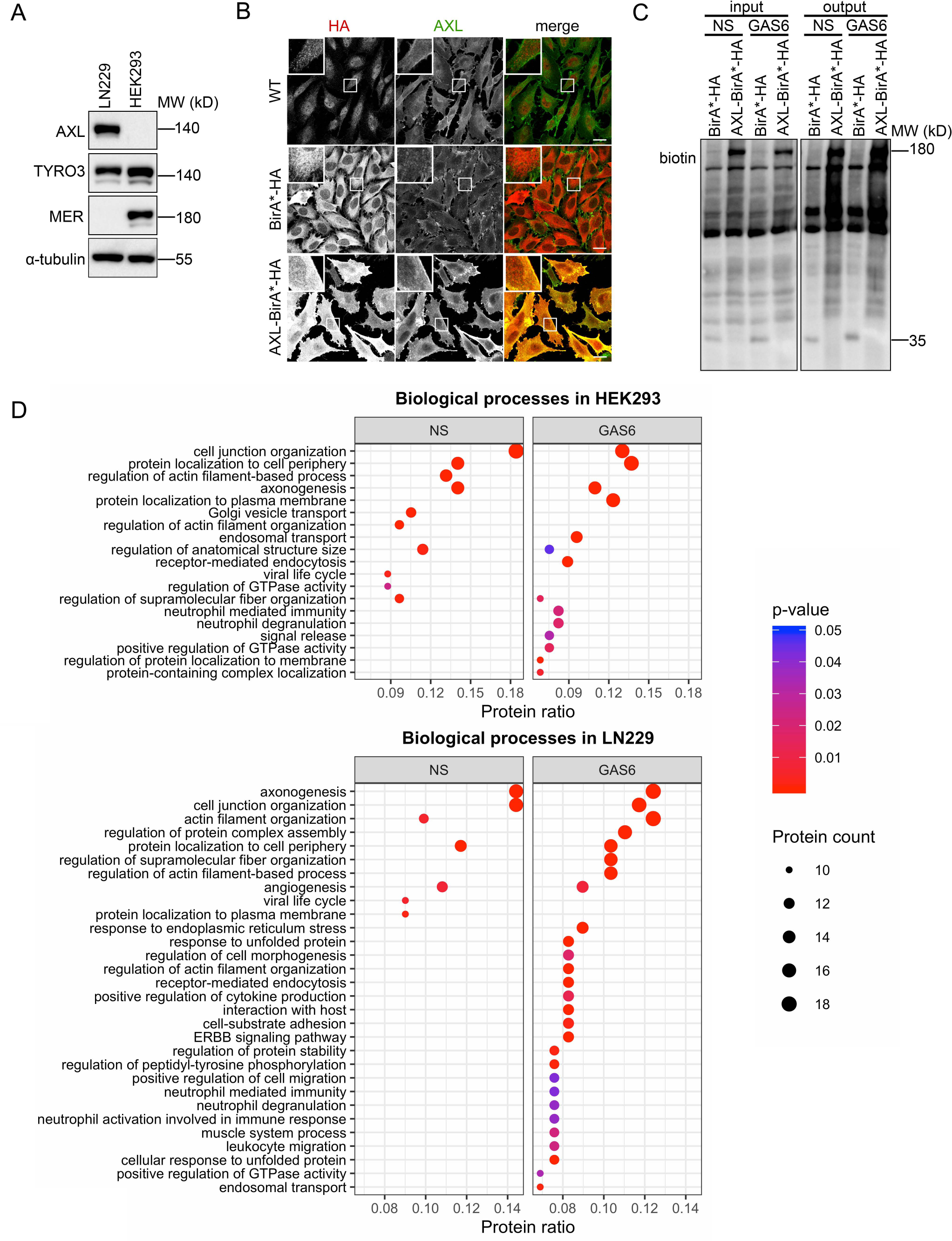
The BioID interactome analysis reveals that AXL is implicated in the regulation of actin cytoskeleton. **A** Western blot showing expression of TAM receptors (AXL, TYRO3, MER) in LN229 and HEK293 cells. α-tubulin was used as a loading control. **B** Confocal images showing the localization of BirA*-HA and AXL-BirA*-HA in LN229 cells stained with antibodies recognizing HA (red) and AXL (green). WT-wild type LN229 cells. **C** Western blot showing the biotinylation status of proteins before (input) and after (output) pull-down with streptavidin-coated magnetic beads. Serum-starved LN229 cells expressing BirA*-HA or AXL-BirA*-HA were incubated with biotin for 24 h in the presence or absence of GAS6, lysed and processed as described in the Methods section. Antibodies against biotin were used for blotting. **D** Graphs showing the GO analysis of biological processes for the BioID hits identified in HEK293 (top panel) and LN229 cells (bottom panel). Data information: Insets in confocal images are magnified views of boxed regions in the main images. Scale bars: 20 μm. NS-non-stimulated cells, GAS6-GAS6-stimulated cells.

The lists of proteins identified in cells expressing AXL-BirA*-HA protein (+/-GAS6) were compared to the ones found in control BirA*-HA expressing cells. Proteins fulfilling the following criteria were considered as AXL proximity interactors: they were identified in at least 2 out of 3 experiments, with ≥ 2 peptides at least in one experiment, and had three times higher Mascot score in AXL-BirA*-HA samples in comparison to the control. We found 116 such proteins in non-stimulated and 151 proteins in GAS6-treated HEK293 cells, whereas we identified 114 and 147 proteins in non-stimulated and GAS6-treated LN229 cells, respectively (Data EV1 and 2). Among interactors identified in HEK293 cells, 26 and 61 proteins were unique for non-stimulated and GAS6-stimulated conditions, respectively, whereas 90 proteins were common for both groups (Fig EV1G). In case of hits identified in LN229 cells, 21 proteins were uniquely found in non-stimulated samples, 54 proteins were unique for GAS6-stimulated samples, and 93 proteins were common (Fig EV1G). The comparison of hits from HEK293 and LN229 cells revealed 56 proteins identified in both lines under non-stimulated conditions, and 67 proteins upon GAS6 stimulation (Fig EV1H).

The Gene Ontology (GO) analysis of biological processes among the identified hits indicated enrichment of proteins implicated in cell junction organization, several actin-related processes, axonogenesis, supramolecular fiber organization and angiogenesis in both non-stimulated and GAS6-stimulated samples. In GAS6-stimulated cells, AXL was found to interact also with proteins involved in receptor-mediated endocytosis and endosomal transport, positive regulation of GTPase activity, signaling, cell-substrate adhesion and positive regulation of migration. Importantly, more proteins implicated in actin-related processes were found in GAS6-treated than in non-stimulated LN229 cells (Fig 1D). The GO analysis of molecular functions also showed enrichment of actin-binding proteins among AXL proximity interactors (Fig EV2A and B). Similarly, the GO analysis of cellular components identified proteins localizing to the cell leading edge and lamellipodium, which are actin-rich structures (Fig EV2C and D).

Generally, the composition of the interactome of AXL, particularly upon GAS6 stimulation, points to the involvement of this receptor in the regulation of actin dynamics as one of its main functions in the cell.

### GAS6 triggers the formation of peripheral and circular dorsal ruffles

Taking into account the importance of actin-related processes for cancer cell migration and invasion (Yamaguchi & Condeelis, 2007), we focused on a possible role of GAS6-AXL signaling in actin cytoskeleton remodeling. First, we visualized actin in LN229 cells using phalloidin staining. We found that AXL accumulated in cell regions enriched in actin, including lamellipodia, both in non-stimulated and GAS6-treated cells (Fig EV3A). Moreover, stimulation of LN229 cells with GAS6 triggered membrane ruffling, manifested by the formation of PRs and CDRs (Fig 2A). CDR formation upon GAS6 stimulation was also confirmed in lung and breast cancer cell lines, A549 and MDA-MB-231, respectively (Fig EV3B and C). Quantification of the number of CDRs in LN229 cells stimulated with GAS6 for different time periods revealed that the maximal CDR formation occurred after 10 min of stimulation and subsided afterwards (Fig 2B and C).

**Fig 2.**
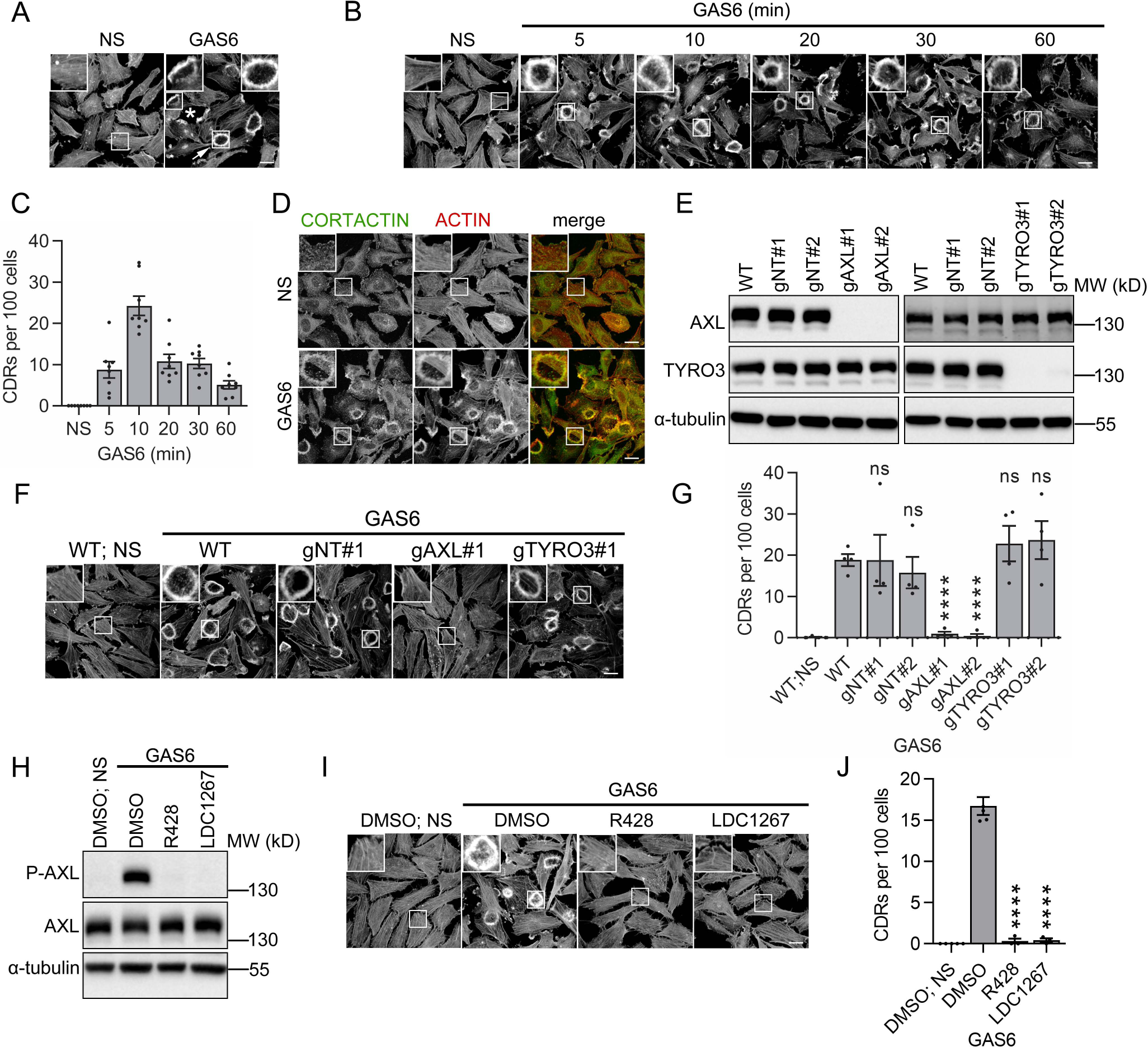
GAS6 induces the formation of CDRs which depends on AXL and its kinase activity. **A** Confocal images showing the formation of PRs (marked with asterisk) and CDRs (marked with arrow). Serum-starved LN229 were stimulated with GAS6 for 10 min, fixed and actin was stained with phalloidin. **B** Confocal images showing the kinetics of CDR formation in LN229 cells stimulated with GAS6 for the indicated time periods. Actin was stained with phalloidin. **C** Quantitation of the CDRs shown in (B), n=8. **D** Confocal images showing cortactin accumulation on CDRs upon stimulation of serum-starved LN229 cells with GAS6 for 10 min. Fixed cells were stained with antibodies recognizing cortactin (green) and phalloidin to visualize actin (red). **E** Western blot showing the efficiency of CRISPR-Cas9-mediated knockout of *AXL* and *TYRO3* in LN229 cells. Two gRNAs targeting *AXL* (gAXL#1 and gAXL#2) and targeting *TYRO3* (gTYRO3#1 and TYRO3#2) were used. CRISPR-Cas9-edited LN229 cells with two non-targeting gRNAs (gNT#1 and gNT#2) served as controls. WT-wild type LN229 cells. α-tubulin served as a loading control. **F** Confocal images showing CDR formation upon knockout of *AXL* and *TYRO3* in LN229 cells (described in E). Serum-starved cells were stimulated with GAS6 for 10 min, fixed and actin was stained with phalloidin. **G** Quantification of data shown in F, n=4. Student’s unpaired t-test, ****p≤0.0001, ns-non-significant (p>0.05). **H** Western blot showing AXL phosphorylation (P-AXL, Y702) upon AXL inhibitors. Serum-starved LN229 cells were pretreated for 30 min with R428 or LDC1267 followed by stimulation with GAS6 for 10 min. α-tubulin served as a loading control. **I** Confocal images showing CDR formation upon AXL inhibitors. Serum-starved LN229 cells were treated as described in (H). Actin was stained with phalloidin. **J** Quantification of data shown in (I), n≥3, Student’s unpaired t-test, ****p≤ 0.0001. Data information: Insets in confocal images are magnified views of boxed regions in the main images. Scale bars: 20 μm. For data quantification approximately 150 cells were counted per experiment, and each dot represents data from one independent experiment whereas bars represent the means ± SEM from n experiments. NS-non-stimulated cells, GAS6-GAS6-stimulated cells.

In line with the observed GAS6-induced CDR formation, a more profound analysis of AXL proximity interactors revealed 23 proteins known to localize and/or contribute to the formation of CDRs (Table EV1). One of them was cortactin, a cytoplasmic actin-binding protein that is considered as one of main CDR markers (Krueger et al, 2003). Immunostaining of GAS6-stimulated LN229 cells confirmed that cortactin colocalized with actin on CDRs (Fig 2D). Cumulatively, these data show that AXL is localized to actin-rich cell regions and GAS6 stimulation rapidly induces the formation of cortactin-positive CDRs.

### GAS6-induced CDR formation depends on AXL and its kinase activity

GAS6 was described as a ligand for all three TAM receptors (Lew et al, 2014; Mark et al, 1996; Nagata et al, 1996; Stitt et al, 1995; Tsou et al, 2014). Since LN229 cells express two of them, AXL and TYRO3 (Fig 1A), we assessed the contribution of these receptors to GAS6-induced CDR formation. To this end, we knocked out *AXL* and *TYRO3* by the CRISPR-Cas9 method and tested GAS6-induced CDR formation in the generated knockout LN229 lines. Western blot analyses confirmed efficient inactivation of *AXL* or *TYRO3* with two different gRNAs per gene (Fig 2E). We discovered that the knockout of AXL, but not TYRO3, inhibited GAS6-driven CDRs (Fig 2F and G). The same effects were observed when expression of *AXL* and *TYRO3* was silenced by siRNAs (Fig EV3D-F).

To determine whether activation of the intracellular tyrosine kinase domain of AXL was required for the formation of GAS6-induced CDRs, we tested the effect of AXL inhibitors on the appearance of these actin structures. Treatment of LN229 cells with R428 or LCD1267 for 30 min prior to stimulation with GAS6 prevented AXL phosphorylation and CDR formation (Fig 2H-J). Altogether, these data indicate that the formation of CDRs upon GAS6 stimulation depends on AXL and its kinase activity.

### PI3K mediates GAS6-induced CDR formation downstream of the GAS6-AXL signaling pathway

AXL has been shown to trigger activation of several downstream signaling pathways, such as PI3K-AKT, ERK, and PLC-γ (Paccez et al, 2014). However, activation of a particular set of downstream effectors is often cell type-specific. To verify which proteins are activated by GAS6-AXL signaling and can mediate GAS6-induced CDR formation in glioblastoma cells, we assessed the phosphorylation status of different effectors in lysates of GAS6-stimulated LN229 cells using a phospho*-*kinase array (Fig 3A, Fig EV4A). As shown in Fig 3A, GAS6 mainly triggered phosphorylation of AKT and its substrates, such as GSK3αβ, PRAS40 and WNK1 (Nascimento et al, 2006). This potent activation of AKT was also confirmed by Western blot (Fig 3B and C), but its pharmacological inhibition with MK-2206 had no effect on GAS6-induced CDR formation (Fig 3D and E, Fig EV4B). AKT phosphorylation indicated that GAS6 might activate PI3K, thus we checked whether pharmacological inhibition of PI3K activity affected GAS6-stimulated CDRs. Two inhibitors specific for class I PI3Ks, GDC-0941 and ZSTK474, prevented the GAS6-AXL-dependent formation of CDRs in LN229 cells (Fig 3F and G, Fig EV4C). These results are in agreement with a previous report that PDGF-induced CDRs were blocked by inhibition of PI3K but not AKT (Corallino et al, 2018).

**Fig 3.**
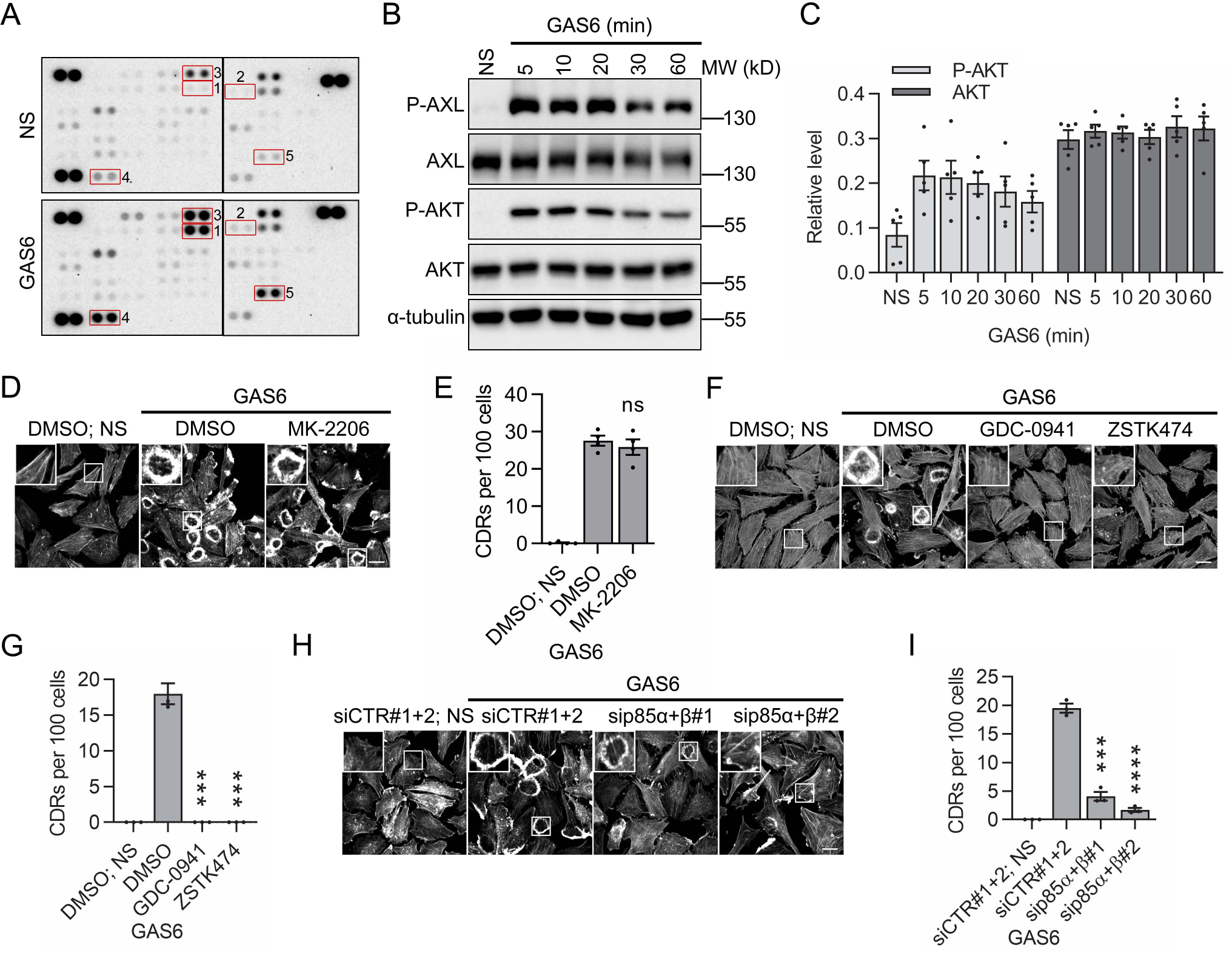
PI3K mediates GAS6-induced CDR formation. **A** A phospho*-*kinase array showing activation of AKT and its substrates in serum-starved LN229 cells upon stimulation with GAS6 for 10 min. 1. AKT (S473) 2. AKT (T308) 3. GSK3αβ (S21/S9) 4. PRAS40 (T246) 5. WNK1 (T60). Three paired spots in the corners are positive controls. **B** Western blot showing GAS6-induced phosphorylation of AXL (P-AXL, Y702) and AKT (P-AKT, S473). Serum-starved LN229 cells were stimulated with GAS6 for the indicated time periods. α-tubulin served as a loading control. **C** Graph showing the densitometric analysis of P-AKT and AKT levels shown in (B), n=5. The levels of P-AKT and AKT were normalized to α-tubulin. **D** Confocal images showing GAS6-triggered CDR formation upon AKT inhibitor. Serum-starved LN229 cells were pretreated with MK-2206 for 30 min prior to stimulation with GAS6 for 10 min, fixed and actin was stained with phalloidin. **E** Quantification of CDR formation shown in (D), n=4. Student’s unpaired t-test, ns-non-significant (p>0.05). **F** Confocal images showing CDR formation in GAS6-stimulated cells pretreated with PI3K inhibitors. Serum-starved LN229 cells were incubated with GDC-0941 or ZSTK474 for 30 min followed by incubation with GAS6 for 10 min. Next, cells were fixed and actin was stained with phalloidin. **G** Quantification of CDR formation shown in (F), n=3. Student’s unpaired t-test, ***p≤0.001. **F** Confocal images showing formation of GAS6-induced CDRs upon siRNA-mediated depletion of p85 isoforms. LN229 cells were transfected with combination of non-targeting siRNAs (siCTR#1+siCTR#2) or two combinations of siRNAs targeting p85α and p85β (sip85α#1+p85β#1 or sip85α#2+p85β#2). 72 h after transfection serum-starved cells were stimulated with GAS6 for 10 min, fixed and actin was stained with phalloidin. **I** Quantification of CDRs shown in (H), n=3. Student’s unpaired t-test, ***p≤0.001, ****p≤0.0001. Data information: Insets in confocal images are magnified views of boxed regions in the main images. Scale bars: 20 μm. For quantification of microscopic data approximately 150 cells were counted per experiment. Each dot represents data from one independent experiment whereas bars represent the means ± SEM from n experiments. NS-non-stimulated cells, GAS6-GAS6-stimulated cells.

We found two regulatory subunits of class IA PI3Ks, p85α and p85β, among AXL interactors only in GAS6-stimulated samples which indicated that they were specifically associated with activated AXL (Table EV1, Data EV1 and 2). Both subunits have been recently described as downstream effectors of RAB35, and their knockout prevented the formation of PDGF-induced CDRs (Corallino et al, 2018). Thus, to investigate their involvement in GAS6-AXL-driven CDR formation we depleted both p85 isoforms by siRNA-mediated silencing in LN229 cells. Knockdown of p85α and β inhibited CDR formation upon GAS6 stimulation (Fig 3H and I, Fig EV4D). Cumulatively, these findings indicate that AXL stimulation by GAS6 induces mainly PI3K-AKT signaling, whereas GAS6-triggered formation of CDRs is specifically mediated by PI3K.

### GAS6-induced PRs and CDRs drive macropinocytosis through which GAS6-AXL complexes are internalized

It is well established that membrane ruffling promotes macropinocytosis, and both PRs and CDRs were postulated to initiate macropinosome formation (Buckley & King, 2017; Dowrick et al, 1993; Hoon et al, 2012). Given this, we checked whether GAS6-induced membrane ruffling is associated with the formation of macropinosomes. To this end, we performed live imaging of GAS6-stimulated LN229 cells expressing a plasma membrane tethered mCherry (MyrPalm-mCherry). Upon GAS6 stimulation, we observed intense membrane ruffling and formation of large vesicles reminiscent of macropinosomes (Fig 4A, Movie EV1 and 2). We next asked whether GAS6-AXL complexes were internalized via the formed macropinosomes. Confocal imaging of cells immunostained for AXL and Myc-tagged GAS6 revealed that both, the receptor and its ligand, were present on the macropinosome membrane where they colocalized with EEA1 and Rabankyrin-5 (Rank-5), markers of early endosomes and macropinosomes (Schnatwinkel et al, 2004), respectively (Fig 4B and C). We further confirmed that AXL-positive macropinosomes were formed also in other cancer cell lines such as A549, MDA-MB-231 and ovarian cancer SKOV3 cells (Fig EV5A-C). Altogether, these data indicate that GAS6 triggers membrane ruffling that drives internalization of GAS6-AXL complexes via macropinocytosis.

**Fig 4.**
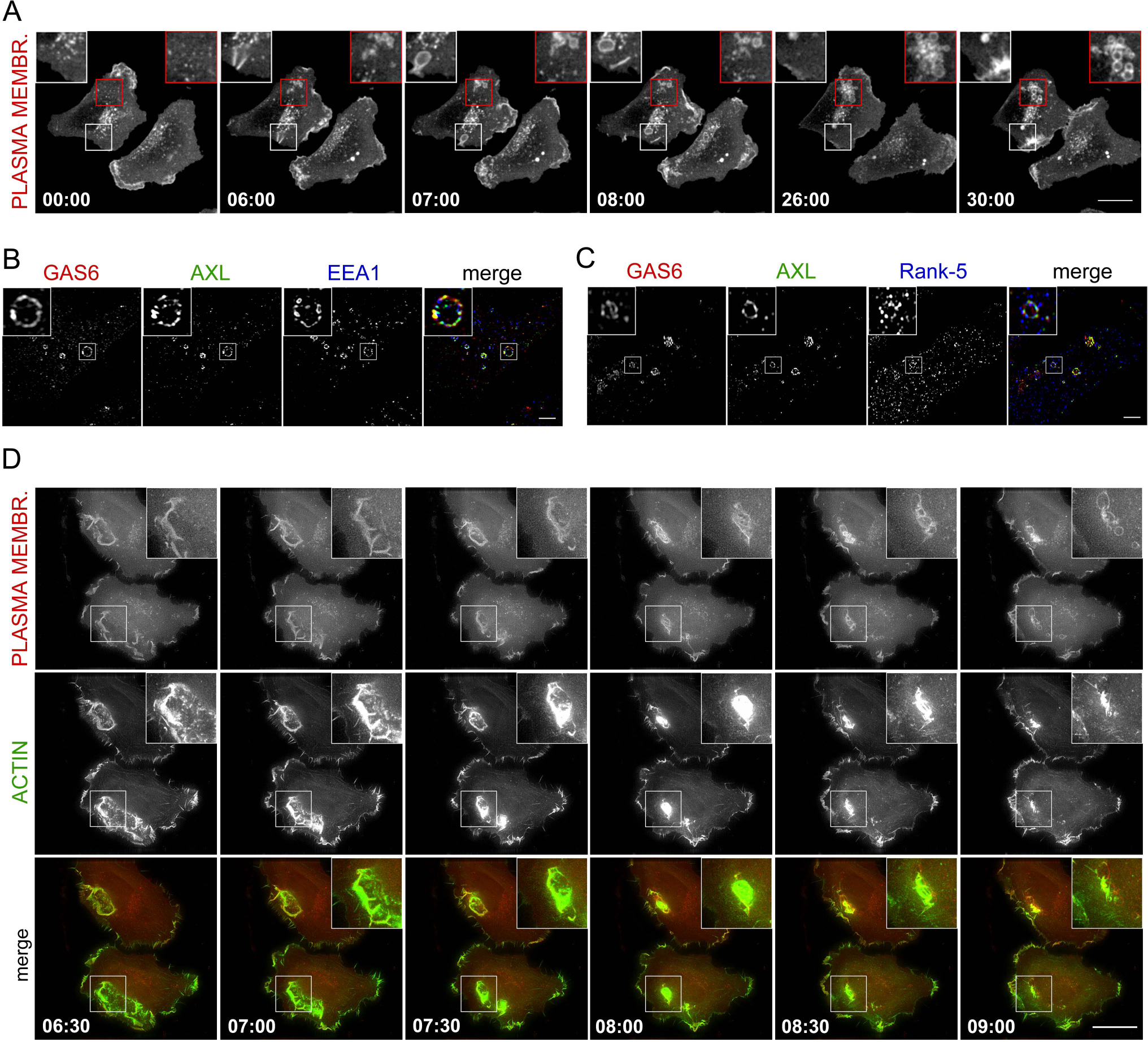
GAS6-induced PRs and CDRs drive macropinocytosis through which GAS6-AXL complexes are internalized. **A** Live-cell time-lapse images of LN229 cells showing the formation of macropinosomes upon GAS6 stimulation. Serum-starved cells expressing the plasma membrane tethered mCherry (MyrPalm-mCherry) were imaged for 5 min before and 30 min after addition of GAS6 by spinning disk confocal microscopy. Representative frames (0-30 min of GAS6 stimulation) from Movie EV1 are shown. Scale bar: 20 μm. **B, C** High resolution confocal images of GAS6-stimulated LN229 cells showing GAS6 and AXL on macropinosomes marked by EEA1 (B) and Rank-5 (C). Serum-starved cells were stimulated with GAS6 for 10 min, fixed and stained with antibodies against AXL (green), Myc (red, to detect GAS6-MycHis) and EEA1 (blue, B) or Rank-5 (blue, C). Scale bars: 2 μm. **D** Live-cell time-lapse images of LN229 cells showing GAS6-induced CDRs and macropinosomes formed upon the closure of CDRs. Serum-starved cells expressing MyrPalm-mCherry (to stain the plasma membrane, red) and LifeAct-mNeonGreen (to stain actin, green) were imaged for 5 min before and 20 min after addition of GAS6 by widefield fluorescence microscopy. Representative frames from Movie EV3 (6:30-9 min of GAS6 stimulation) are shown. Scale bar: 20 μm. Data information: Insets are magnified views of boxed regions in the main images.

Although it has been reported that macropinosomes can be formed upon closure of CDRs, some findings suggest that PRs, but not CDRs, are necessary and sufficient for macropinocytosis (Suetsugu et al, 2003). Thus, to test the involvement of GAS6-induced CDRs in macropinosome formation, we performed live imaging of GAS6-stimulated LN229 cells expressing MyrPalm-mCherry to label the plasma membrane, together with LifeAct-mNeonGreen that stained filamentous actin (F-actin). This dual-color live cell imaging confirmed that CDRs formed upon GAS6 stimulation were large, dynamic and transient actin structures (Fig 4D, Movie EV3 and 4). It further revealed that some macropinosomes were indeed formed upon closure of CDRs (Fig 4D, Movie EV3 and 4). However, although GAS6 stimulation triggered appearance of AXL-positive macropinosomes in GAS6-stimulated SKOV3, we did not observe the formation of CDRs in these cells (Fig EV5C).

### GAS6-induced macropinocytic internalization depends on AXL and its downstream activation of PI3K

To determine whether GAS6 stimulates macropinocytic internalization of a model cargo, we measured uptake of a high molecular mass dextran. Immunofluorescence analysis showed that fluorescently labeled dextran was internalized into large vesicular structures in LN229 cells within 10 min of GAS6 stimulation (Fig 5A). A quantitative analysis of the acquired images showed an almost twofold increase in integral fluorescence intensity of dextran-positive vesicles in GAS6-stimulated cells in comparison to non-stimulated cells (Fig 5B). In parallel, measurements of dextran uptake by flow cytometry confirmed increase in dextran internalization upon GAS6 stimulation (Fig 5C-E, Fig EV6). GAS6-induced dextran uptake was completely inhibited by a macropinocytosis inhibitor 5-(N-ethyl-N-isopropyl)amiloride (EIPA) (Koivusalo et al, 2010) (Fig 5C, Fig EV6B). Next, we checked whether kinase activity of AXL was needed for the observed GAS6-stimulated macropinocytic internalization of dextran. As shown in Fig 5C and Fig EV6B, incubation of LN229 cells with AXL inhibitors, R428 and LDC1267, prior to GAS6 stimulation completely blocked GAS6-induced uptake of dextran. This indicates that activation of the kinase domain of AXL is required for GAS6-driven macropinocytosis, similarly to GAS6-induced CDR formation. Moreover, siRNA-mediated depletion of AXL, but not of TYRO3, completely inhibited GAS6-stimulated macropinocytic dextran internalization (Fig 5D, Fig EV6C and D). These data clearly demonstrate that, although LN229 cells express two TAM receptors, AXL and TYRO3, GAS6-activated macropinocytic uptake of dextran depends specifically on AXL. PI3K-generated PI(3,4,5)P_3_ has been shown to play an important role during macropinocytosis (Hoeller et al, 2013; Rupper et al, 2001). Since PI3K is the main downstream signaling pathway activated by GAS6-AXL in LN229 cells, we assessed the impact of pharmacological inhibition of PI3K on GAS6-stimulated macropinocytic dextran internalization. We found that the incubation of LN229 cells with PI3K inhibitors, GDC-0941 and ZSTK474, prior to GAS6 stimulation blocked GAS6-induced uptake of dextran (Fig 5E, Fig EV6E). Cumulatively, these findings indicate that, similarly to CDR formation, GAS6-induced macropinocytosis depends on AXL and its kinase activity as well as downstream activation of PI3K but not on a related TAM receptor TYRO3.

**Fig 5.**
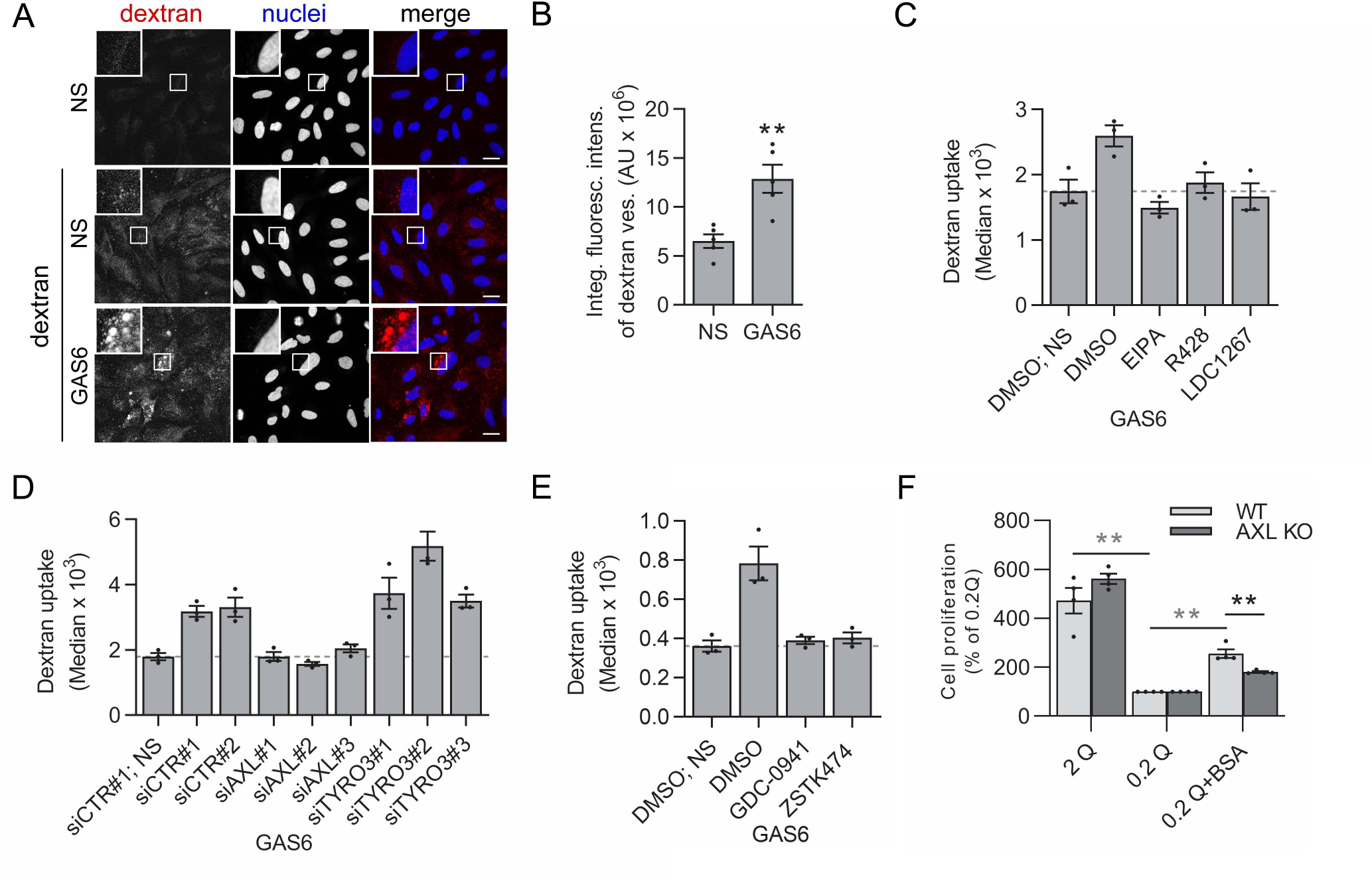
GAS6 induces macropinocytic internalization of dextran which depends on AXL and downstream PI3K activation. **A** Confocal images showing internalization of fluorescently labeled dextran (red) into large vesicular structures following stimulation with GAS6. Serum-starved LN229 cells were incubated with GAS6 and dextran for 10 min, fixed and stained with DAPI to visualize nuclei (blue). **B** Analysis of integral fluorescence intensity of dextran-positive vesicles (expressed in arbitrary units, AU) shown in (A), n=5. Student’s unpaired t-test **p≤0.01 **C-E** Graphs showing GAS6-induced macropinocytic uptake of dextran upon macropinocytosis (EIPA) and AXL inhibitors (R428 and LDC1267) (C), siRNA-mediated depletion of AXL or TYRO3 (D) and PI3K inhibitors (GDC-0941 and ZSTK474) (E) in LN229 cells. For experiments with inhibitors, serum-starved cells were incubated for 30 min with the indicated inhibitor prior to GAS6 stimulation. For siRNA-mediated depletion two non-targeting siRNAs (siCTR#1 and siCTR#2), three siRNAs against AXL (siAXL#1, siAXL#2, siAXL#3) and three siRNAs against TYRO3 (siTYRO3#1, siTYRO3#2, siTYRO3#3) were used. Cells pretreated with inhibitors or cells 72 h post-transfection were stimulated with GAS6 for 2 min, next fluorescent dextran was added, and cells were incubated for additional 20 min. The fluorescence of internalized dextran was measured by flow cytometry, n=3. **F** Graph showing the proliferation of wild type (WT) and AXL knockout (KO) LN229 cells incubated in medium with normal (2 mM Q) or sub-physiological (0.2 mM Q) glutamine concentration in the absence or presence of 2% BSA. AXL KO cells were generated using gAXL#2. The medium was replaced every 24 h and cell proliferation was measured after 6 days using an ATPlite assay. Data are presented relative to the values obtained for LN229 cells grown in 0.2 mM Q. Student’s unpaired or one sample (gray stars) t-test **p≤0.01, ns-non-significant (p>0.05). Data information: Insets in confocal images are magnified views of boxed regions in the main images. Scale bars: 20 μm. For quantification of microscopic data approximately 150 cells were counted per experiment. Each dot represents data from one independent experiment whereas bars represent the means ± SEM from n experiments. Dashed lines show median fluorescence of dextran taken up by non-stimulated cells. NS-non-stimulated cells, GAS6-GAS6-stimulated cells.

### AXL-activated macropinocytosis of albumin partly rescues growth of glioblastoma cells in sub-physiological concentrations of glutamine

It was demonstrated that RAS-transformed pancreatic cancer cells acquire glutamine via the lysosomal degradation of exogenously provided albumin, internalized through macropinocytosis, which allows them to survive and proliferate in the absence of this amino acid (Commisso et al, 2013). Thus, we hypothesized that GAS6-AXL-induced macropinocytosis of albumin could similarly promote survival of glioblastoma LN229 cells. To verify this possibility we examined whether proliferation of LN229 cells was affected by glutamine deprivation, and whether this effect could be reversed by culturing cells in media supplemented with 2% bovine serum albumin (BSA). To avoid growth arrest and to reproduce conditions used previously by others (Commisso et al, 2013), experiments with glutamine deprivation were done in the presence of 10% dialysed serum (in contrast to standard GAS6 stimulations in our study performed in the absence of serum). Since GAS6 is present in serum (Balogh et al, 2005), to estimate the contribution of AXL to cell growth, we tested both wild type and AXL knockout LN229 cells. As shown in Fig 5F, proliferation of LN229 cells was significantly inhibited under sub-physiological glutamine concentration (0.2 mM Q), which was partly rescued by addition of BSA. Moreover, the survival of AXL knockout cells in BSA-supplemented media was lower than the wild type cells, indicating that AXL contributes to the observed albumin-mediated improvement in cell viability (Fig 5F). Thus, our results suggest that AXL-dependent macropinocytosis allows exploiting exogenous albumin to improve survival of LN229 cells under conditions of glutamine deprivation.

### GAS6-AXL-induced CDRs accumulate components of focal adhesions (FAs)

In addition to macropinocytosis, another postulated function of CDRs is remodeling of cellular components in preparation for migration that requires disassembly of adhesion sites. In this regard, integrin β3 and β1 were shown to be rapidly and transiently redistributed to CDRs upon focal adhesion disassembly during cell migration stimulated by growth factors (Gu et al, 2011). Given this, we tested the localization of integrin β1 in GAS6-stimulated LN229 cells which were seeded on fibronectin-coated coverslips. We observed accumulation of integrin β1 on GAS6-induced CDRs (Fig 6A), along with other FA components such as vinculin, talin and paxillin (Fig 6B-D).

**Fig 6.**
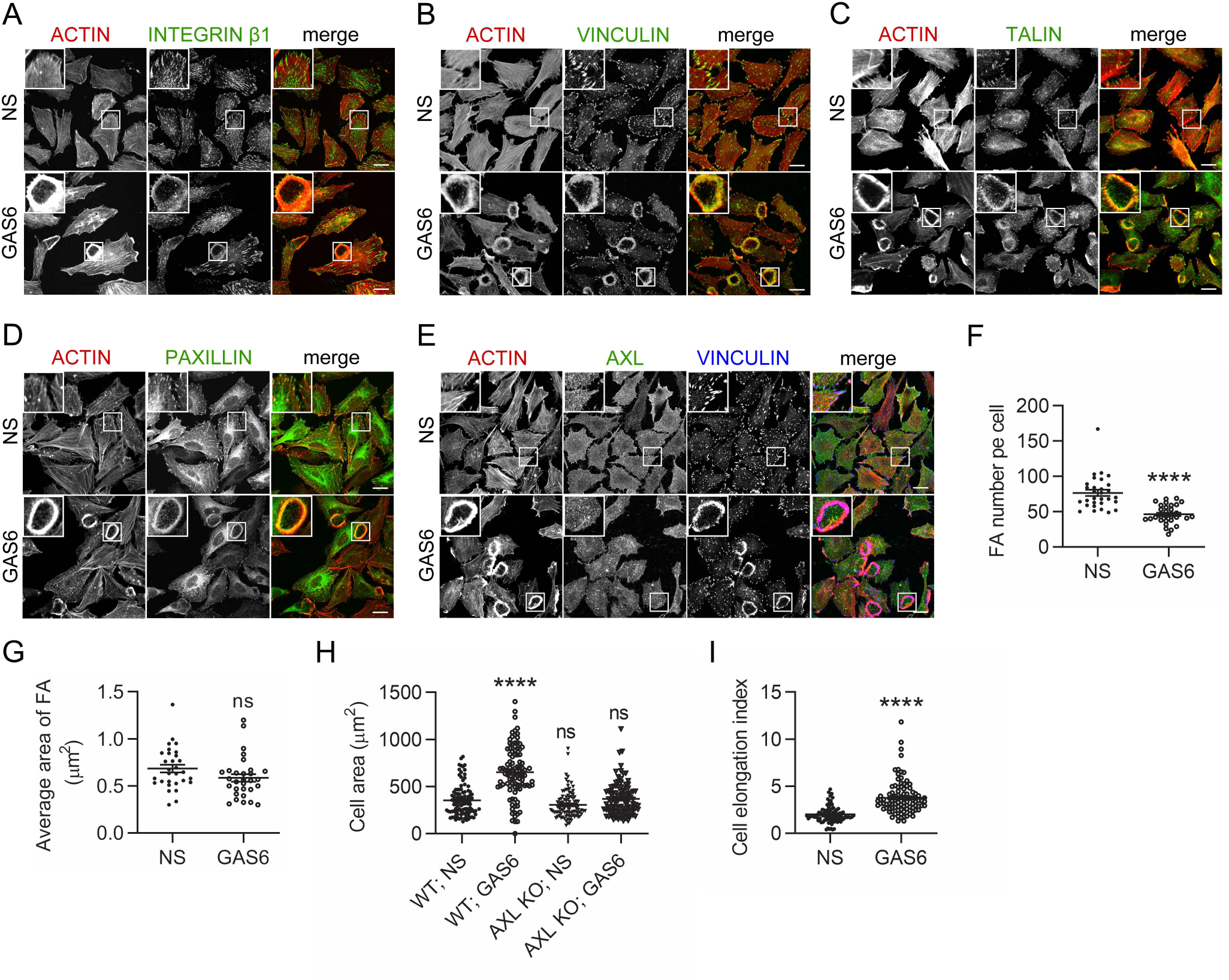
GAS6-AXL-induced CDRs trigger disassembly of FAs via transient accumulation of their components. **A-D** Confocal images of LN229 cells showing accumulation of integrin β1 (A), vinculin (B), talin (C) and paxillin (D) on GAS6-induced CDRs. Serum-starved cells were stimulated with GAS6 for 10 min, fixed and stained with antibodies against integrin β1 (A), vinculin (B), and talin (C) (green). To visualize paxillin, LN229 cells expressing paxillin tagged with mNeonGreen were used (D, green). Actin was stained with phalloidin (red). **E** Confocal images of LN229 cells showing accumulation of vinculin, but not AXL, on GAS6-induced CDRs. Serum-starved cells were stimulated with GAS6 for 10 min, fixed and stained with antibodies against AXL (green) and vinculin (blue). Actin was stained with phalloidin (red). **F, G** Graphs showing the number (F) and average area of FAs (G) in serum-starved LN229 cells upon stimulation with GAS6 for 10 min. FAs and CDRs were visualized by staining of vinculin and actin, respectively. The number and the area of FAs per cell were counted using ImageJ software for 10 non-stimulated and 10 GAS6-stimulated, CDR-containing cells from three independent experiments (n=30). Mann–Whitney U test, ****p≤0.0001, ns-non-significant (p>0.05). **H** Graph showing spreading of wild type (WT), and AXL knockout (KO) LN229 cells following GAS6 stimulation. AXL KO cells were generated using gAXL#2. Cells were plated onto fibronectin-coated coverslips and incubated in the absence or presence of GAS6 for 30 min, fixed and actin was stained with phalloidin. Cell area was measured using ImageJ software for 50 non-stimulated and 50 GAS6-stimulated cells from two independent experiments (n=100). Mann–Whitney U test, ****p≤0.0001, ns-non-significant (p>0.05). **I** Graph showing cell elongation index of serum-starved LN229 cells upon stimulation with GAS6 for 10 min. Fixed cells were stained with phalloidin to visualize actin. The cell elongation index (the ratio between the major axis and the minor axis) was measured using ImageJ software for 50 non-stimulated and 50 GAS6-stimulated, CDR-containing cells from two independent experiments (n=100). Mann–Whitney U test, ****p≤0.0001. Data information: Insets in confocal images are magnified views of boxed regions in the main images. Scale bars: 20 μm. NS-non-stimulated cells, GAS6-GAS6-stimulated cells. Each data point represents a value for a single cell, horizontal lines are means ± SEM for n cells.

Our GO analysis of cellular components among the identified AXL interactors revealed enrichment of proteins localized to FAs, including these FA components which were accumulated on GAS6-induced CDRs (Fig EV2C and D, Data EV1 and 2). Despite the biochemical detection of many FA components in close proximity to AXL in the BioID assay, we did not observe evident accumulation of AXL on FAs independently of the presence or absence of serum (Fig 6E, Fig EV7A). Similarly, we did not detect AXL on CDRs formed upon 10 min stimulation with GAS6 (Fig 6E). However, we observed accumulation of AXL together with vinculin on lamellipodia, which are sites of new FA formation (Zaidel-Bar et al, 2004) (Fig EV7A). Altogether, these findings revealed that GAS6-driven CDRs concentrate FA components suggesting that they may be involved in turnover of existing FAs.

### GAS6-AXL-induced CDRs are involved in FA disassembly

To test whether GAS6-induced CDRs affected FA dynamics we quantified the number and area of these adhesion structures in non-stimulated and GAS6-stimulated cells exhibiting CDRs. As shown in Fig 6F and EV7B, GAS6-stimulated LN229 cells with CDRs contained considerably fewer FAs in comparison to non-stimulated cells. In contrast, average FA area was not changed between non-stimulated and GAS6-treated cells with CDRs (Fig 6G). These observations suggested that GAS6-induced CDRs might lead to FA disassembly through transient accumulation of FA components. Since during attachment to the ECM substrate, cell spreading requires continuous formation and disassembly of FAs (Huveneers & Danen, 2009) we assessed the impact of 30 min GAS6 stimulation on LN229 cells plated on fibronectin. As shown in Fig 6H, GAS6-stimulated LN229 cells spread better in comparison to non-stimulated cells. Moreover, this GAS6-dependent increase in cell spreading was reduced in AXL knockout cells (Fig 6H, Fig EV7C).

Finally, we calculated a cell elongation index (the ratio between the major and minor axis of cell, Fig EV7D) which directly correlates with mesenchymal migration, a specific mode of cell locomotion through the ECM, associated with CDR-dependent matrix invasion (Hoon et al, 2012; Suetsugu et al, 2003; Zobel et al, 2018). We found that the cell elongation index of GAS6-stimulated cells forming CDRs was significantly higher in comparison to non-stimulated cells (Fig 6I, Fig EV7D). Cumulatively, these observations allow concluding that GAS6-AXL-induced CDRs are implicated in FA turnover by concentration of FA components. They further suggest that CDRs might play a role in cell migration through the ECM.

### The GAS6-AXL signaling pathway induces cell invasion in a PI3K-dependent manner

RAB35-dependent CDRs were proposed to act as a steering wheel for PDGF- and HGF-mediated chemotaxis and chemoinvasion (Corallino et al, 2018). To check whether GAS6 is a chemotactic factor, we tested migration of LN229 cells through a membrane towards GAS6 gradient in a Boyden chamber. We found that, in contrast to serum, GAS6 did not attract LN229 cells (Fig EV8A). Since it was postulated that CDRs might correspond to invasive protrusions and play a role during 3D invasion (Hoon et al, 2012; Sero et al, 2011; Suetsugu et al, 2003), we tested the impact of GAS6 on invasion of spheroids formed by cancer cells. As shown in Fig 7A and B, GAS6 stimulated invasion of LN229 spheroids into Matrigel and the same effect was also observed for MDA-MB-231 spheroids (Fig EV8B and C). Similarly to CDR formation and macropinocytosis, GAS6-stimulated invasion was inhibited by the CRISPR-Cas9-mediated inactivation of AXL but not of TYRO3 (Fig 7A and B). Moreover, invasion of LN229 spheroids was dependent on the kinase activity of AXL, as it was completely blocked by two AXL inhibitors, R428 and LDC1267 (Fig 7C and D).

**Fig 7.**
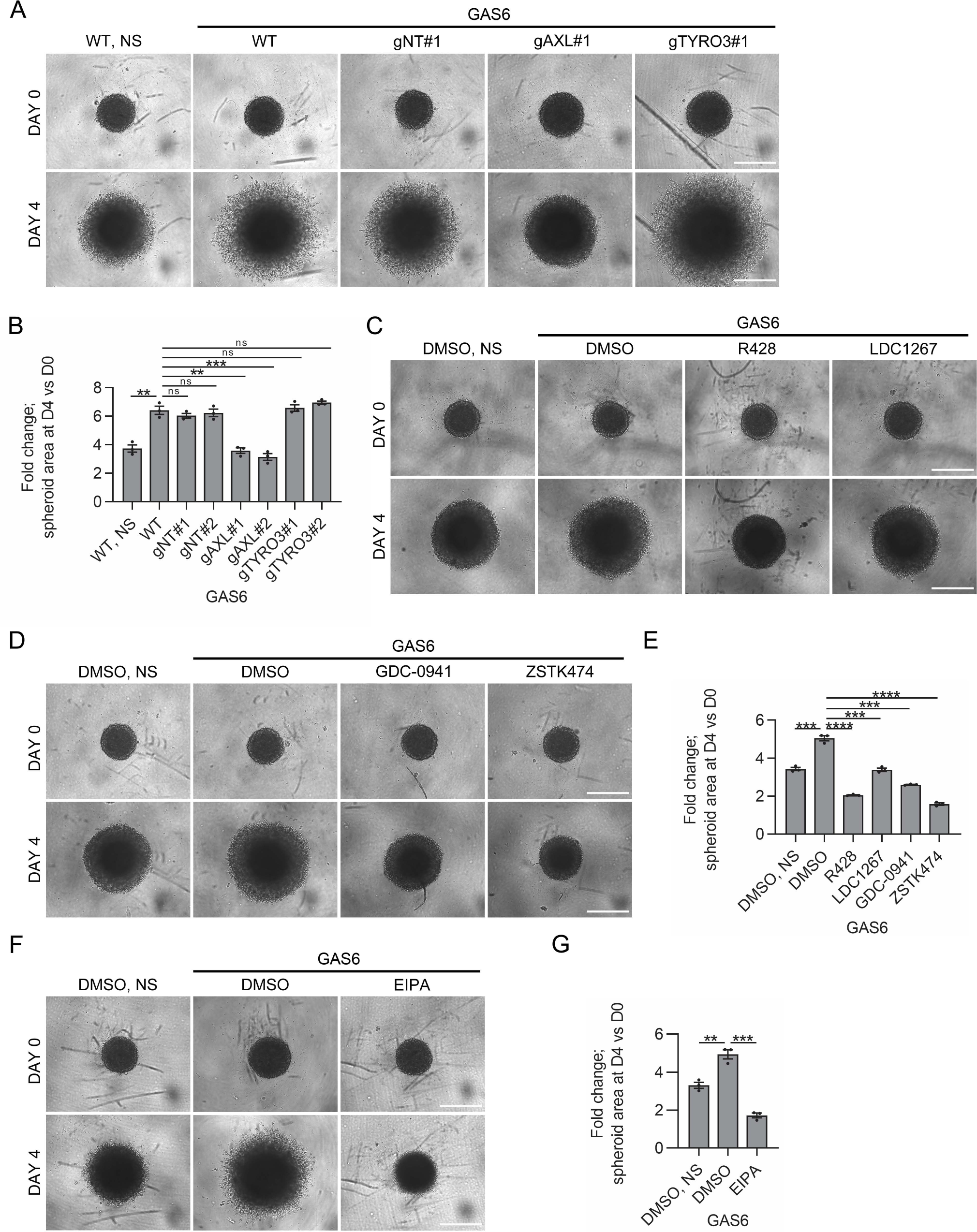
GAS6-AXL signaling pathway induces cell invasion in a PI3K-dependent manner. **A** Images showing GAS6-stimulated invasion of wild type (WT) and CRISPR-Cas9-mediated knockout of *AXL* and *TYRO3* LN229 cells. Two different gRNAs targeting *AXL* (gAXL#1 and gAXL#2) and targeting *TYRO3* (gTYRO3#1 and TYRO3#2) were used. CRISPR-Cas9-edited LN229 cells with two different non-targeting gRNAs (gNT#1 and gNT#2) served as controls. Spheroids were grown in the presence or absence of GAS6 in Matrigel for 4 days. Scale bars: 500 μm. **B** Quantification of data shown in (A), n=3. Student’s unpaired t-test, **p≤0.01, ***p≤0.001, ns-non-significant (p>0.05). **C, D, F** Images showing GAS6-induced invasion of LN229 spheroids after treatment with inhibitors of AXL (R428 or LDC1267) (C), PI3K (GDC-0941 and ZSTK474) (D) and macropinocytosis (EIPA) (F). Spheroids embedded in Matrigel were incubated with the indicated inhibitor for 30 min followed by incubation with GAS6 for 4 days. Scale bars: 500 μm. **E, G** Quantification of data shown in (C, D) and (F), respectively, n=3. Student’s unpaired t-test, ***p≤0.001, ****p≤0.0001, ns-non-significant (p>0.05). Data information: The area of spheroids was measured by ImageJ software. Data are expressed as fold changes of the spheroid area on the 4^th^ day (DAY 4, D4) with respect to the spheroid area before Matrigel addition (DAY 0, D0). Each dot represents data from one independent experiment whereas bars represent the means ± SEM from n experiments. NS-non-stimulated cells, GAS6-GAS6-stimulated cells.

Since a pharmacological inhibition of PI3K blocked both formation of CDRs and macropinocytosis following GAS6 stimulation (Fig 3F and G, Fig 5E), we assessed whether PI3K inhibitors prevented GAS6-induced invasion of LN229 spheroids. As shown in Fig 7D and E, PI3K inhibitors, GDC-0941 and ZSTK474, completely blocked GAS6-induced invasion of LN229 spheroids into Matrigel. GAS6-stimulated invasion was also blocked by the macropinocytosis inhibitor EIPA (Fig 7F and G). This suggests that macropinocytosis may be involved in GAS6-AXL-induced invasion of cancer cells.

Collectively, these findings show that GAS6 induces cancer cell invasion which depends on AXL, the activity of its tyrosine kinase domain and downstream activation of PI3K but not on its cognate TAM receptor TYRO3. Moreover, our data indicate that GAS6-induced CDRs and macropinocytosis may contribute to invasion triggered by the activation of GAS6-AXL signaling.

## Discussion

AXL, a member of the TAM receptor family, and its ligand GAS6 are associated with pathogenesis and metastasis of multiple cancers (Paccez et al, 2014; Verma et al, 2011; Wu et al, 2014). Despite active development of AXL inhibitors for clinical use in oncology, astonishingly little is known about its intracellular mechanisms of action. Here, we identified the interactome of AXL in HEK293 and glioblastoma LN229 cells and thereby found that GAS6-AXL signaling regulates several actin-related processes that enhance an invasive phenotype of cancer cells. GAS6 triggered intense membrane ruffling, in the form of PRs and CDRs, and induced macropinocytosis. Furthermore, GAS6-induced CDRs promoted the turnover of FAs via accumulation of their components. Functionally, AXL activation by GAS6 culminated in increased invasion of tumor spheroids. At the molecular level, we found that PI3K-AKT was the main signaling pathway downstream of GAS6-AXL activation. A pharmacological inhibition of PI3K or depletion of its regulatory subunits, p85α and p85β, blocked CDR formation, macropinocytic internalization and invasion of spheroids induced by GAS6. Based on our results we propose that GAS6-AXL signaling induces PI3K-dependent, actin-driven cytoskeletal rearrangements and macropinocytosis that jointly contribute to cancer cell invasion (Fig 8).

**Fig 8.**
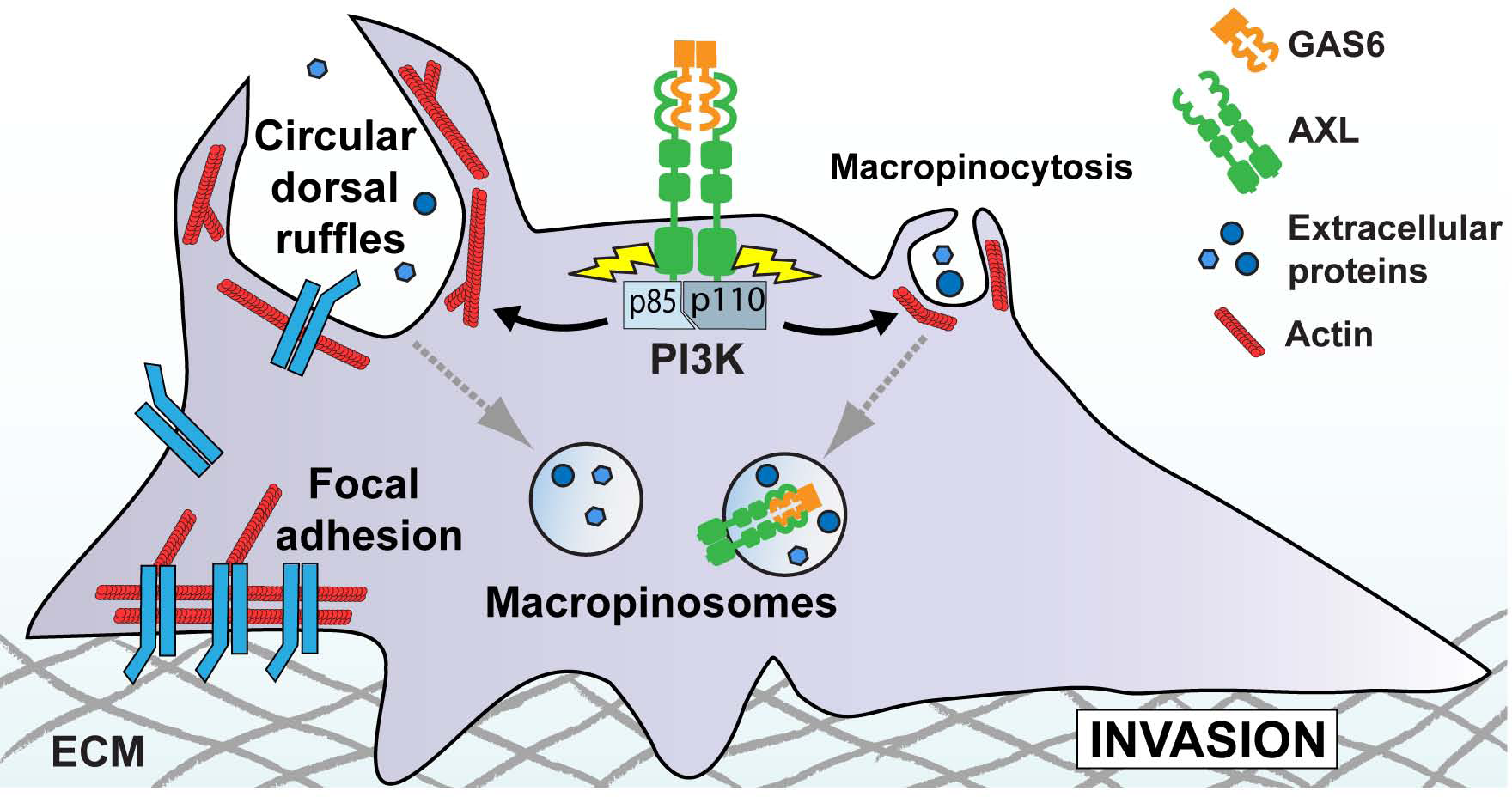
GAS6-AXL signaling induces actin remodeling that drives membrane ruffling, macropinocytosis, FA disassembly and cancer cell invasion. Upon GAS6 binding AXL associates with two regulatory subunits of class IA PI3Ks, p85α and p85β, that activates PI3K signaling. Then, GAS6-AXL-PI3K signaling triggers membrane ruffling, both PRs and CDRs, that drives internalization of GAS6-AXL complexes via macropinocytosis. Macropinocytic uptake of extracellular proteins improves growth of cancer cells in a nutrient-poor environment. In parallel, GAS6-induced CDRs contribute to the disassembly of FAs through recruitment of their components, thus preparing cells for mesenchymal migration. Altogether, activation of GAS6-AXL-PI3K signaling induces cancer cell invasion where GAS6-induced CDRs and macropinocytosis contribute to this process.

There are two known ligands for the TAM receptors, GAS6 and PROS1. However, the specificity and biological relevance of their interaction with TAMs are not fully elucidated. It has been shown that, in addition to AXL, GAS6 can also activate two other TAMs, TYRO3 and MER (Lew et al, 2014; Tsou et al, 2014). Here, we discovered that depletion of AXL or pharmacological inhibition of its kinase activity completely inhibited GAS6-induced membrane ruffling, macropinocytosis and invasion of LN229 cells grown as spheroids. In contrast, TYRO3, another TAM receptor expressed by LN229 cells, was dispensable for all these processes. Thus, our findings unequivocally demonstrate that AXL is a primary receptor for GAS6, at least in the glioblastoma model used.

Upon ligand binding, RTKs undergo endocytosis, via different, often parallel, internalization pathways (Goh & Sorkin, 2013). Endocytosis is an important regulator of cellular signaling activated by RTKs that, if aberrant, contributes to RTK-associated tumorigenesis (Miaczynska, 2013). However, AXL endocytosis has not been systematically investigated. Here, we discovered that by inducing actin remodeling GAS6 stimulation triggers macropinosome formation, and GAS6-AXL complexes localize to macropinosomes indicating that a fraction of AXL is internalized via macropinocytosis. Thus, our data provide a mechanistic explanation for previous virology studies reporting that AXL might contribute to macropinocytic uptake of Lassa and Ebola viruses upon infection (Fedeli et al, 2018; Hunt et al, 2011).

There are different possible mechanisms by which macropinocytosis could contribute to GAS6-activated intracellular trafficking of AXL. For example, Chiasson-MacKenzie et al (2018) demonstrated that the deficiency of merlin, a cytoskeletal ERM-like protein, induced CDR formation and macropinocytosis that favored internalization of EGFR and its subsequent recycling. We found merlin among AXL proximity interactors. Thus, it is conceivable that GAS6-induced CDRs and macropinocytosis might depend on AXL-mediated phosphorylation of merlin that inhibits its function. A report indicated that CDRs may also constitute an endocytic mechanism of EGFR internalization (Orth et al, 2006). Here, we did not observe GAS6 or AXL accumulation on CDRs formed upon 10 min stimulation with GAS6. This indicates that, in contrast to EGFR, AXL does not undergo CDR-driven internalization. However, it is possible that the observed accumulation of AXL and actin on the dorsal site of cells following 5 min stimulation with GAS6 reflects an early stage of CDR formation. This could suggest that AXL may be recruited to the site of CDR formation at the very beginning and displaced from CDRs at later stages of their formation. Altogether, our study provides foundational data about ligand-induced endocytic trafficking of AXL, a process which has not been previously studied for any of the TAM receptors.

Macropinosomes have been postulated to form concomitantly with the contraction and closure of CDRs (Hoon et al, 2012). However, Suetsugu et al (2003) questioned the involvement of these structures in macropinosome formation. Here, by time-lapse live imaging of GAS6-stimulated LN229 cells expressing markers of the plasma membrane and actin, we revealed that macropinosomes were indeed generated upon closure of CDRs. However, the formation of macropinosomes following GAS6 stimulation was not accompanied by CDR formation in SKOV3 cells. Thus, we postulate that PR- and CDR-derived macropinosomes may represent separate populations of macropinosomes, and their molecular composition and function may differ. An existence of different types of macropinosomes with diverse functions is an exciting possibility that merits further research.

Cancer metastasis remains a major clinical problem. The driving mechanisms of each step of the metastatic cascade are still poorly characterized due to their complexity. Therefore, defining such mechanisms and their key molecular players is essential, especially to develop more effective therapies (Chitty et al, 2018; Menezes et al, 2016; Steeg, 2016). A large body of evidence documents that AXL is an important regulator of cancer progression, invasion and metastasis in a plethora of malignant tumors (Kimani et al, 2016; Paccez et al, 2014; Verma et al, 2011; Wu et al, 2014). Thus, to identify biological processes and novel molecular players underlying an AXL-dependent aggressive phenotype we established its interaction network using the BioID assay. To our knowledge, a comprehensive AXL interactome has not been reported so far. The GO analysis of the identified AXL interactors indicated, among others, enrichment of proteins implicated in actin dynamics. Accordingly, we found that AXL was accumulated in actin-rich cell regions, including lamellipodia, in glioblastoma LN229 cells. Similarly, a very recent study of Zajac *et al*. (2020) showed that AXL colocalized with F-actin at the leading edge of migrating mesenchymal triple-negative breast cancer cells, and its depletion impaired the directionality of cell migration (Zajac et al, 2020).

To leave a primary tumor and invade into the surrounding ECM and tissues, cancer cells have to migrate through complex 3D environments. It requires active remodeling of actin cytoskeleton and formation of specialized actin-rich structures and protrusions. For this reason, many actin-associated proteins, also those identified here as AXL interactors like cortactin or WAVE2, are frequently overexpressed in metastatic cancers (Yamaguchi & Condeelis, 2007). Here, we describe several actin-dependent processes downstream of AXL activation in cancer cells, such as CDR formation, FA turnover and macropinocytosis. Together with literature data, these findings reveal that the GAS6-AXL signaling axis drives multiple cellular processes involving actin remodeling that jointly result in an invasive phenotype. Moreover, the interaction network of AXL including numerous actin-associated proteins may explain its previously postulated contribution to EMT that could result from controlling actin-dependent processes needed for epithelial cells to acquire a mesenchymal phenotype (Sun et al, 2015; Yilmaz & Christofori, 2009).

Here, we discovered that GAS6 stimulation triggered the formation of AXL-dependent CDRs, actin structures implicated in invasion. Previous studies and our data showed that CDRs are linked to FA turnover, as proteins from disassembling FAs are redistributed and transiently accumulated on CDRs (Gu et al, 2011; Reinecke et al, 2014). We identified many FA components as AXL proximity interactors that independently confirms the role of AXL in the regulation of FA dynamics. AXL may also contribute to the formation of new FAs on the cell leading edge since it was accumulated together with vinculin on lamellipodia, that are sites of new FA assembly. Since mesenchymal migration of cancer cells requires constant disassembly of existing FAs and assembly of new ones, our results indicate that the GAS6-AXL signaling pathway may regulate these processes, partially by triggering the formation of CDRs. AXL was recently shown to stimulate the formation and activity of invadopodia in melanoma cells (Revach et al, 2019). This is another type of actin-based protrusions important for cancer metastasis, as they degrade and remodel the ECM (Eddy et al, 2017). Invadopodia and FAs share many components (Wang & McNiven, 2012), some of which are AXL proximity interactors. This, together with the fact that we observed AXL accumulation on cortactin- and actin-positive dot-like structures in LN229 cells (data not shown), may indicate that this receptor is concentrated on invadopodia.

Macropinocytosis is another process discovered here that may contribute to AXL-promoted cancer progression and invasion. Several studies demonstrated that macropinocytosis, through uptake of extracellular proteins (e.g. albumin) and their further degradation in lysosomes, enables cancer cells to survive in a nutrient-poor tumor microenvironment (Commisso et al, 2013; Lee et al, 2019; Palm, 2019; Recouvreux & Commisso, 2017). However, the majority of these studies concentrated on macropinocytosis triggered by mutated RAS and/or were conducted in pancreatic cancer cells, whereas data on RTK-mediated macropinocytosis as an alternative nutrient uptake route in other cancer types were lacking. Here, we showed that AXL-driven macropinocytosis may be exploited by glioblastoma cells for uptake of extracellular albumin to allow them to proliferate under glutamine-poor conditions. Moreover, GAS6-stimulated invasion of LN229 spheroids was blocked upon pharmacological inhibition of macropinocytosis. It is possible that GAS6-induced macropinocytosis promotes also internalization of ECM components, such as fragments of collagen, as shown for pancreatic ductal adenocarcinoma (Olivares et al, 2017). Such macropinocytic uptake of degraded ECM fragments may contribute to cancer invasiveness not only by promoting cell survival under nutrient-limited conditions but also by clearing a path through the ECM for invading tumor cells. Moreover, a very recent study of Jayashankar and Edinger (2020) showed that macropinocytic internalization of necrotic cell debris drives proliferation and drug resistance of breast and prostate cancer with oncogenic mutations that activate KRAS or PI3K. In phagocytes, TAM receptors participate in efferocytosis and TAM ligands are postulated to act as a bridge between phagocyte receptors and phosphatidylserine on apoptotic cells or cell debris (Doran et al, 2019; Lemke & Burstyn-Cohen, 2010; Nguyen et al, 2013). As phagocytosis is mechanistically related to macropinocytosis, it is tempting to speculate that drug resistance of cancers with overactivated GAS6-AXL signaling results, at least partially, from their increased capability for macropinocytic internalization of cell debris.

Finally, our findings demonstrate that PI3K is an important downstream effector in the GAS6-AXL signaling cascade regulating invasion of glioblastoma spheroids via activation of several actin-dependent processes. PI3K was previously implicated in AXL-driven invasion of ovarian cancers (Rankin et al, 2010). These and our own findings suggest that PI3K inhibitors alone or in combination with AXL inhibitors may inhibit metastatic spread of malignant tumors with highly activated GAS6-AXL pathway. Importantly, recent studies demonstrated that the isoform β of PI3K specifically contributes to invasion of breast cancer cells through the regulation of invadopodia maturation (Erami et al, 2019). Since usage of isoform-specific inhibitors can limit the toxicity of panPI3K inhibitors (Hanker et al, 2019) it will be important to test which isoform of PI3K plays a predominant role in AXL-driven invasion. Of note, FDA recently approved Alpelisib (BYL719) as the first PI3K inhibitor, specific for the α isoform, for the treatment of advanced and metastatic breast cancer (FDA, May 24, 2019).

In summary, our study revealed that GAS6-activated AXL triggers cancer cell invasion via induction of multiple actin-dependent processes that impact cell dynamics, adhesion and metabolism. Moreover, AXL interactors identified here may constitute novel molecular targets for the development of effective therapies for patients with advanced and metastatic cancers.

## Materials and Methods

### Reagents

#### Inhibitors

PI3K inhibitors: GDC-0941 (HY-50094, used at 5 µM) and ZSTK474 (HY-50847, used at 5 µM), AXL inhibitors: R428 (HY-15150, used at 5 µM) and LDC1267 (HY-12494, used at 5 µM), AKT inhibitor: MK-2206 (HY-10358, used at 0.5 µM) all from MedChemExpress; macropinocytosis inhibitor 5-(N-Ethyl-N-isopropyl)amiloride (EIPA) from Sigma-Aldrich (A3085, used at 25 µM). **Other:** vitamin K_1_ (3804.1, Carl Roth GmbH), puromycin (Toku-E, P001, used at 1 µg/mL), Geneticin® Selective Antibiotic (G418, 11811031, used at 1 mg/mL), Dextran, Tetramethylrhodamine, 70,000 MW (D1818, used at 1 mg/mL), Dextran, Texas Red™, 70,000 MW (D1864, used at 0.25 mg/mL), Dextran, Oregon Green™ 488, 70,000 MW (D7173, used at 0.25 mg/mL) all from Thermo Fisher Scientific; fibronectin (F1056, used at 20 μg/mL), Bovine Serum Albumin (A1470), DAPI (D9542), phalloidin-Atto 390 (50556) all from Sigma-Aldrich.

### Antibodies

The following primary antibodies were used: rabbit anti-HA (sc-805, Western blot (WB) 1:1000), goat anti-AXL (sc-1096, immunofluorescence (IF) 1:500), rabbit anti-AXL (sc-20741, WB 1:1000), mouse anti-c-Myc (sc-40, IF 1:500) all from Santa Cruz Biotechnology; goat anti-AXL (AF154, IF 1:500, WB 1:1000) from R&D Systems; mouse anti-AKT (2920, WB 1:2000), rabbit anti-phospho-AKT (Ser 473) (4060, WB 1:1000), rabbit anti-TYRO3 (5585, WB 1:1000), rabbit anti-phospho-AXL (Tyr702) (5724, WB 1:1000) all from Cell Signaling Technology; mouse anti-α-tubulin (T5168, WB 1:10,000), mouse anti-vinculin (V9131, IF 1:500), mouse anti-cortactin (05-180, IF 1:400), mouse anti-talin (T3287, IF 1:200) all from Sigma-Aldrich; rabbit anti-EEA1 (ALX-210-239, IF 1:1000) from Enzo Life Sciences; mouse anti-EEA1 (610457, IF 1:1000) from BD Biosciences; rabbit anti-biotin (ab53494, WB 1:1000), mouse anti-integrin beta 1 (ab30394, IF 1:400) all from Abcam; rabbit anti-Rabankyrin-5 was a gift from Professor Marino Zerial (Max Planck Institute of Molecular Cell Biology and Genetics, Dresden, Germany (Schnatwinkel et al, 2004), IF 1:500).

Secondary antibodies used for WB: horseradish peroxidase (HRP)-conjugated anti-mouse-IgG (111-035-062), anti-rabbit-IgG (111-035-144) and anti-goat-IgG (805-035-180) antibodies were from Jackson ImmunoResearch; anti-rabbit-IgG conjugated to IRDye 680 (926-68023) and anti-mouse-IgG conjugated to IRDye 800CW (926-32212) antibodies used in the Odyssey system were from LICOR Biosciences. Secondary antibodies used for IF: Alexa Fluor 488-, 555-, 647-conjugated anti-goat-IgG, anti-mouse-IgG and anti-rabbit-IgG were from Thermo Fisher Scientific.

### Plasmids

Plasmid encoding human GAS6, pSecTag2-hGAS6, was a gift from Professor Raymond B. Birge (Rutgers, New Jersey Medical School, Newark, New Jersey, USA (Tsou et al, 2014)). To construct a plasmid encoding GAS6 tagged with Myc-His (pSecTag2-hGAS6-MycHis) the stop codon from pSecTag2-hGAS6 was removed using the QuickChange II XL Site-Directed Mutagenesis kit (Agilent Technologies). Next, to obtain pcDNA3.1-GAS6-MycHis, a sequence encoding the secretion signal from the V-J2-C region of the mouse Ig kappa-chain with hGAS6 was subcloned from pSecTag2-hGAS6-MycHis into pcDNA3.1^(+)^ Mammalian Expression Vector (Thermo Fisher Scientific) using NheI/PmeI restriction sites.

pMyrPalm-mCherry was described elsewhere (Schink et al, 2017). pLifeAct-mNeonGreen was generated by exchanging the mCherry fluorophore in pLifeAct-mCherry (Schink et al, 2017) to mNeonGreen. pLV-pEF1α-mNeonGreenPXN was generated by cloning of a sequence encoding human paxillin (PXN) from pmCherry Paxillin (a gift from Kenneth Yamada, Addgene plasmid #50526; http://n2t.net/addgene:50526; RRID:Addgene_50526) using forward 5’-TGTTCTCTCGAGCATGGACGACCTCGACGCCCTG-3’ and reverse 5’-GTGCTTGCGGCCGCCTAGCAGAAGAGCTTGAGGAAG-3’ oligonucleotides (containing XhoI and NotI restriction sites, respectively) into pLV-pEF1α-mNeonGreen vector (a gift from Dr. Franz Meitinger, Ludwig Institute for Cancer Research, San Diego, USA).

LentiCRISPRv2 was a gift from Feng Zhang (Addgene plasmid #52961; http://n2t.net/addgene:52961; RRID:Addgene_52961). Lentiviral packaging plasmids: psPAX2 (a gift from Didier Trono, Addgene plasmid # 12260; http://n2t.net/addgene:12260; RRID:Addgene_12260) and pMD2.G (a gift from Didier Trono, Addgene plasmid # 12259; http://n2t.net/addgene:12259; RRID:Addgene_12259). LentiCRISPRv2 plasmids with non-targeting gRNAs were a gift from Dr. K. Mleczko-Sanecka (International Institute of Molecular and Cell Biology in Warsaw, Poland). gRNA sequences for CRISPR-Cas9-mediated gene inactivation (Table EV2) were cloned into the LentiCRISPRv2 vectors using a protocol described elsewhere (Sanjana et al, 2014).

To obtain pcDNA3.1-AXL-BirA*-HA plasmid, AXL coding sequence was amplified from pcDNA3.1-AXL using forward 5’ TGTTCTGCTAGCATGGCGTGGCGGTGCCCCAG-3’ and reverse 5’-GTGCTTGGATCCGGCACCATCCTCCTGCCCTG-3’ oligonucleotides (containing NheI and BamHI restriction sites, respectively) and subcloned into pcDNA3.1-MCSBirA(R118G)-HA vector (a gift from Kyle Roux, Addgene plasmid #36047; http://n2t.net/addgene:36047; RRID:Addgene_36047, (Roux et al, 2012)). The pcDNA3.1-AXL was constructed by amplification of cDNA encoding human AXL from human CCD-1070Sk fibroblasts using forward 5’-TGTTCTCTTAAGATGGCGTGGCGGTGCCCCAG-3’ and reverse 5’-GTGCTTCTCGAGTCAGGCACCATCCTCCTGCC-3’ oligonucleotides (containing AlfII and XhoI restriction sites, respectively) and subcloning into pcDNA3.1^(+)^.

To obtain pLenti-CMV-MCS-BirA*-HA, BirA*-HA was amplified from pcDNA3.1-MCSBirA(R118G)-HA using forward 5’-TGTTCTTCTAGAGCTAGCGCTTAAGGCCTGTTAAC-3’ and reverse 5’-GTGCTTACGCGTCTATGCGTAATCCGGTACATC-3’ oligonucleotides (containing XbaI and MluI restriction sites, respectively) and subcloned into pLenti-CMV-MCS-GFP-SV-puro (a gift from Paul Odgren, Addgene plasmid #73582; http://n2t.net/addgene:73582; RRID:Addgene_73582, (Witwicka et al, 2015)). Similarly, to obtain pLenti-CMV-MCS-AXL-BirA*-HA, AXL-BirA*-HA was amplified from pcDNA3.1-AXL-BirA*-HA using forward 5’ TGTTCTGCTAGCATGGCGTGGCGGTGCCCCAG-3’ and reverse 5’-GTGCTTACGCGTCTATGCGTAATCCGGTACATC-3’ oligonucleotides (containing NheI and MluI restriction sites, respectively) and subcloned into pLenti-CMV-MCS-GFP-SV-puro.

### Cell culture

LN229, MDA-MB-231, SKOV3, HEK293, HEK293T and CCD-1070Sk cells were purchased from ATCC, and A549 cells were purchased from Sigma-Aldrich. LN229, MDA-MB-231, A549, HEK293 and HEK293T cells were maintained in DMEM, SKOV3 cells were cultured in McCoy’s 5a medium and CCD-1070Sk cells in MEM. The media were supplemented with 10% fetal bovine serum (FBS) and 2 mM L-glutamine (all from Sigma-Aldrich). Cells were cultured at 37 °C and 5% CO_2_ and regularly tested for mycoplasma contamination.

### Generation of stable HEK293 cell line secreting GAS6-MycHis and purification of GAS6-MycHis from conditioned medium

HEK293 cells were transfected with pcDNA3.1-GAS6-MycHis linearized with PvuI restriction enzyme using Lipofectamine® 2000 Transfection Reagent (Thermo Fisher Scientific), according to the manufacturer’s protocol, and stable transfectants were selected by culturing in medium supplemented with G418 for 2 weeks. Next, single clones were isolated and secretion of GAS6-MycHis into medium was verified by Western blot. To obtain single clones, 500 cells were seeded on 10 cm dish in full medium and cultured for 2 weeks. Once colonies were formed, the medium was removed from the dish and colonies were moved to 24-well plate by scratching cells with sterile 200 μL pipette tips. The selected HEK293 clone expressing high levels of recombinant GAS6 was cultured for 5 days in serum-free medium, supplemented with 10 µg/mL vitamin K_1_ to produce conditioned medium. The medium with GAS6-MycHis was next dialyzed against 50 mM phosphate buffer pH 7.4, with 300 mM NaCl and 10 mM imidazole, and GAS6-MycHis was purified on HisPur Cobalt Resin (Thermo Fisher Scientific) according to the manufacturer’s instructions. Protein purity and yield were assessed by 10% SDS-PAGE followed by Coomassie Brilliant Blue (R-250) staining of the gel. Fractions containing GAS6-MycHis were pooled and dialyzed against PBS using Amicon Ultra-15 Centrifugal Filter Unit (Millipore). The concentration of purified GAS6-MycHis (denoted as GAS6) was measured with Pierce™ BCA Protein Assay Kit (Thermo Fisher Scientific).

### Cell stimulation with GAS6 and treatment with inhibitors

LN229 cells were seeded either directly in 6-well plates for WB (typically 3.5×10^5^ cells/well or 2×10^5^ cells/well for siRNA transfection experiments), or on 12-mm coverslips in 24-well plates for IF (typically 5×10^4^ cells/well or 2×10^4^ cells/well for siRNA transfection experiments or 2.5×10^4^ cells/well for measurement of the cell elongation index). Before GAS6 stimulation, cells were incubated in serum-free medium for 16 h. On the day of stimulation, 1 M HEPES pH 7.5 was added to final concentration of 20 mM, and cells were next incubated with 400 ng/mL GAS6 for 10 min at 37 °C outside of the CO_2_ incubator, unless indicated otherwise. For inhibitor treatment, cells were incubated with appropriate concentration of the indicated inhibitor for 30 min at 37 °C prior to stimulation with GAS6. In control samples, the same volume of DMSO was added. In case of siRNA transfection experiments, cells were stimulated with GAS6 72 h after transfection.

### Immunofluorescence (IF) staining and image analysis

After stimulation with GAS6, cells were washed twice with ice-cold PBS for 5 min and fixed with 3.6% paraformaldehyde in PBS for 10 min at room temperature, followed by staining according to the immunofluorescence protocol with saponin permeabilization described elsewhere (Sadowski et al, 2013). For staining shown in Fig 6A and EV7A, cells were permeabilized in 0.1% Triton X-100 in PBS for 10 min, washed twice with PBS for 5 min, and free aldehyde groups were quenched via incubation in 0.5 mL of 50 mM NH_4_Cl for 15 min. Next, cells were washed twice with PBS for 5 min, blocked in 10% FBS in PBS for 30 min and incubated with appropriate primary antibodies for 1 h, followed by 30 min incubation with appropriate fluorescent secondary antibodies. The antibodies were diluted in 5% FBS in PBS. Phalloidin-Atto 390 or DAPI to stain actin or nuclei, respectively, were added during incubation with fluorescent secondary antibodies.

Twelve-bit images with resolution 1024×1024 pixels were acquired using the LSM 710 confocal microscope (Zeiss) with ECPlan-Neofluar 40×1.3 NA oil immersion objective. ZEN 2 software (Zeiss) was used for acquisition. At least ten images were acquired per experimental condition. In case of data shown in Fig 4B and C, confocal laser scanning microscope ZEISS LSM 800 equipped with Plan-Apochromat 63×/1.40 NA oil objective and the Airyscan detection unit (Zeiss) was employed as described elsewhere (Sadowski et al, 2013). Pictures were assembled in Photoshop (Adobe) with only linear adjustments of contrast and brightness. The number of CDRs and FAs, area of FAs and cells, as well as the cell elongation index were counted manually using ImageJ software. Dextran internalization was analyzed by the MotionTracking software (http://motiontracking.mpi-cbg.de). The intracellular accumulation of dextran was expressed as integral fluorescence of all detectable dextran-positive vesicles (Collinet et al, 2010; Rink et al, 2005).

### Generation of HEK293 cells stably expressing BirA*-HA or AXL-BirA*-HA

To generate HEK293 cells expressing BirA*-HA or AXL-BirA*-HA, cells were transfected with pcDNA3.1-BirA*-HA or pcDNA3.1-AXL-BirA*-HA plasmids linearized with PvuI, using Lipofectamine® 2000 Transfection Reagent (Thermo Fisher Scientific), according to the manufacturer’s protocol. Next, cells were grown for 2 days and then selected by culturing in medium supplemented with G418. Single clones were obtained as described above. Expression of BirA*-HA or AXL-BirA*-HA was checked using WB and IF.

### Generation of modified LN229 cells (expressing BirA*-HA, AXL-BirA*-HA and mNeonGreen-paxillin or bearing CRISPR-Cas9 mediated AXL-and TYRO3-knockout)

Cell lines were established via lentiviral transduction of LN229 cells, as described elsewhere (Banach-Orlowska et al, 2018). Lentiviral particles were produced in HEK293T cells using psPAX2 and pMD2.G packaging plasmids and an appropriate plasmid, pLenti-CMV-MCS-BirA*-HA for expression of BirA*-HA, pLenti-CMV-MCS-AXL-BirA*-HA for expression of AXL with the C-terminal BirA*-HA, pLV-pEF1α-mNeonGreenPXN for expression of paxillin tagged with the N-terminal mNeonGreen, or LentiCRISPRv2 with cloned gRNA for the generation of CRISPR-Cas9-mediated knockout. To establish knockout cell lines, two gRNAs sequences targeting *AXL* or *TYRO3,* from the Brunello library, were used (Table EV2) (Doench et al, 2016). Cells stably expressing BirA*-HA or AXL-BirA*-HA and CRISPR-Cas9-mediated knockout lines were selected with puromycin, whereas LN229 cells overexpressing mNeonGreenPXN were selected with G418. Finally, expression or knockout were confirmed by WB and/or IF.

### Proximity-dependent protein identification (BioID) assay

HEK293 cells (1×10^6^) expressing BirA*-HA or AXL-BirA*-HA were seeded on 10 cm dishes in complete medium. Cells were then incubated with 50 μM biotin in the presence or absence of GAS6 for 24 h. LN229 cells (1×10^6^) expressing BirA*-HA or AXL-BirA*-HA were seeded on 10 cm dishes in complete medium. Next day they were starved for 16 h in serum-free medium prior to treatment with 50 μM biotin in the presence or absence of GAS6 in serum-free medium for 24 h. Finally, both cell lines were lysed and biotinylated proteins were isolated using Dynabeads™ MyOne™ Streptavidin C1 (65002, Thermo Fisher Scientific) following the protocol described elsewhere (Roux et al, 2013). Before mass spectrometry analysis, protein biotinylation was verified by WB. The experiments were performed in three biological repeats.

### Mass spectrometry of biotinylated proteins

Mass spectrometry analysis was performed by the Mass Spectrometry Laboratory at Institute of Biochemistry and Biophysics, Polish Academy of Sciences, Warsaw, Poland. Isolated biotinylated proteins were reduced with 5 mM TCEP (for 60 min at 60 °C). To block reduced cysteines, 200 mM MMTS at a final concentration of 10 mM was added, and the samples were incubated at room temperature for 10 min. Next, trypsin (Promega) was added at a 1:20 v/v ratio and samples were incubated at 37 °C overnight. Finally, trifluoroacetic acid was used to inactivate trypsin. Peptide mixtures were analyzed by liquid chromatography coupled to tandem mass spectrometry (LC-MS/MS) using Nano-Acquity (Waters Corporation) UPLC system and LTQ-FT-Orbitrap (Thermo Scientific) mass spectrometer. Measurements were carried out in the positive polarity mode, with capillary voltage set to 2.5 kV. A sample was first applied to the Nano-ACQUITY UPLC Trapping Column using water containing 0.1% formic acid as a mobile phase. Next, the peptide mixture was transferred to Nano-ACQUITY UPLC BEH C18 Column using an acetonitrile gradient (5–35% acetonitrile over 160 min) in the presence of 0.1% formic acid with a flow rate of 250 nL/min. Peptides were eluted directly to the ion source of the mass spectrometer. Each LC run was preceded by a blank run to ensure that there was no carry-over of material from previous analysis. HCD fragmentation was used. Up to 10 MS/MS events were allowed per each MS scan.

Acquired raw data were processed by Mascot Distiller followed by Mascot Search (Matrix Science, London, UK, on-site license) against the SwissProt database restricted to human sequences. Search parameters for precursor and product ions mass tolerance were 30 ppm and 0.1 Da, respectively, enzyme specificity: trypsin, missed cleavage sites allowed: 1, fixed modification of cysteine by methylthio and variable modification of methionine oxidation. Peptides with Mascot score exceeding the threshold value corresponding to <5% expectation value, calculated by Mascot procedure, were considered to be positively identified.

The mass spectrometry proteomics data have been deposited to the ProteomeXchange Consortium via the PRIDE partner repository (Perez-Riverol et al, 2019) with the dataset identifier PXD017933.

### GO analysis of mass spectrometry data

Lists of hits obtained from mass spectrometry analysis of AXL-BirA*-HA samples (with or without GAS6) were compared to the ones obtained from BirA*-HA expressing cells considered as control samples. Proteins identified in at least 2 out of 3 experiments, with ≥ 2 peptides at least in one experiment, and having three times higher Mascot score in AXL-BirA*-HA samples in comparison to control, were considered as AXL proximity interactors. AXL interactors were subjected to the GO analysis of biological processes, molecular functions and cellular components using the clusterProfiler package (version 3.6.0; (Yu et al, 2012)) taking advantage of the enrichGO function. All enrichment p-values in GO analysis were corrected for multiple testing using the Benjamini–Hochberg method, and only proteins with adjusted p-value < 0.05 were considered significant. The minimal and maximal sizes of protein clusters were set to 10 and 500, respectively. Redundant terms were removed by means of the simplify function with cutoff 0.65. Calculations were performed in R version 3.6.1 (https://www.R-project.org).

### siRNA transfection

Twenty-four hours after seeding, LN229 cells were transfected with siRNA using Lipofectamine RNAiMAX (Thermo Fisher Scientific) according to the manufacturer’s protocol. The total concentration of siRNA was 10 nM in case of single depletion or 20 nM if two siRNAs were used. Cells were analyzed 72 h upon transfection and silencing efficiency was controlled by WB or qRT-PCR. Sequences of the used Ambion Silencer Select siRNAs (Thermo Fisher Scientific) are listed in Table EV3.

### Western blot (WB) and quantitative real-time PCR (qRT-PCR)

WB and qRT-PCR were performed as described elsewhere (Banach-Orlowska et al, 2018). Primers for qRT-PCR used to estimate the expression of genes of interest are listed in Table EV4. Data were quantified using Data Assist v2.0 software (Applied Biosystems) and normalized to the level of *ACTB* (actin) mRNA.

### Live-cell imaging

Live-cell imaging of LN229 cells expressing MyrPalm-mCherry was performed using the Opera Phenix spinning disk confocal microscope (Perkin Elmer, Waltham, MA, USA). LN229 cells were seeded on SensoPlate 96-well glass bottom plates from Greiner Bio-One (655892). Images were obtained with 40x/1.1 water immersion objective and Harmony software (version 4.8; Perkin Elemer) was used for image acquisition and analysis. Images were acquired every 30 s starting from 5 min before GAS6 administration till 30 min after addition of the ligand. At least two 16-bit image sequences (with resolution 2048x2048 pixels and binning 2) per experimental condition were acquired. Images were exported to Fiji (version 2.0.0-rc-61/1.52p) to assemble pictures (selected frames) and movies, with only linear adjustment of contrast and brightness.

Live-cell imaging of LN229 cells expressing MyrPalm-mCherry and LifeAct-mNeonGreen was performed using a Deltavision OMX V4 microscope. Images were acquired every 30 s starting from 5 min before addition of GAS6 till 20 min after the ligand administration and were further deconvolved as described elsewhere (Sneeggen et al, 2019).

### Human Phospho-Kinase Array

Phosphorylation status and total amounts of 43 kinases were measured using Human Phospho-Kinase Array from R&D Systems. 1.5×10^6^ LN229 cells were stimulated with 400 ng/mL of GAS6 for 10 min at 37 °C, as described in “Cell stimulation with GAS6” and cells were further processed according to the manufacturer’s protocol.

### Dextran uptake assays

For microscopy-based dextran uptake assay, serum-starved LN229 cells, seeded on 12-mm coverslips in 24-well plates, were stimulated with GAS6 in the presence of 70,000 MW dextran conjugated with tetramethylrhodamine for 10 min at 37 °C, as described in “Cell stimulation with GAS6”. For flow cytometry-based dextran uptake assay, serum-starved LN229 cells, seeded in 6-well plates, were stimulated with GAS6 for 2 min at 37 °C, as described in “Cell stimulation with GAS6”. Next, 70,000 MW dextran conjugated with Texas Red™ (Fig 5C and D) or Oregon Green™ 488 (Fig 5E) was added and cells were incubated for additional 20 min at 37 °C. Finally, cells were washed five times with ice-cold PBS, trypsinized, and fluorescence of internalized dextran was measured by flow cytometry. Data shown in Fig 5C and D were generated using LSRII SORP (BD Biosciences) whereas data shown in Fig 5E were generated using BD LSRFortessa flow cytometer (BD Biosciences).The obtained data were plotted and analyzed by FlowJo (Tree Star Inc.) A total number of 20,000 -50,000 cells were counted for each experimental condition.

### Glutamine deprivation assay

Wild type (WT) and AXL knockout (gAXL#2) LN229 cells were seeded in triplicates into 96-well plates (2×10^3^ cells/well) and cultured in complete medium for 24 h. Next, cells were washed once with PBS and incubated in medium containing 10% dialyzed FBS and sub-physiological glutamine concentration (0.2 mM). Where indicated, 2% BSA was added. The medium was replaced every 24 h. Cells were cultured for 6 days and cell proliferation was measured using ATPlite Luminescence Assay (PerkinElmer).

### Cell spreading assay

Wild type (WT) and AXL knockout (gAXL#2) LN229 cells were trypsinized and serum-starved as described elsewhere (Reinecke et al, 2014). Next, 8×10^4^ cells were seeded in the presence or absence of GAS6 onto glass coverslips coated with 20 µg/mL fibronectin. After 30 min of incubation cells were fixed and actin was stained with phalloidin-Atto 390.

### Transwell migration assay

LN229 cells were seeded into the upper compartment of a 6.5 mm Transwell® with 8.0 µm Pore Polycarbonate Membrane Insert (3422, Corning) (2.5×10^4^ cells/insert) in serum-free medium, whereas the lower chamber contained serum-free medium alone or supplemented with 400 ng/mL GAS6 or 10% FBS. After 20 h incubation at 37 °C cells present on the top of the membrane were removed using cotton swabs, and cells that migrated through the membrane to the lower side were fixed in 70% ethanol and stained with crystal violet.

### Spheroid invasion assay

To form spheroids, 5×10^3^ cells/well cells were seeded into ultra-low attachment 96-well round bottom plate (7007, Corning Costar) and cultured for 4 days in complete DMEM. After visual confirmation of the spheroid formation, 100 µL of medium was removed from each well and replaced with 100 µL of Matrigel® Growth Factor Reduced (GFR) Basement Membrane Matrix (356230, Corning), diluted to the final concentration of 5 mg/mL in ice-cold serum-free DMEM. In order to allow Matrigel to solidify, spheroids were incubated for 1 h at 37 °C and next 100 µL of serum-free DMEM, with or without 3x concentrated GAS6, were added to the final concentration of 400 ng/mL. When relevant, LN229 spheroids were incubated with inhibitors for 30 min prior to addition of GAS6. Images of spheroids were taken before addition of Matrigel (DAY 0) and four days later (DAY 4) using JuLI microscope (Digital Bio). The area of spheroids was measured manually using ImageJ software.

### Statistical methods

Data are provided as means ± SEM from at least three independent experiments, unless stated otherwise. Statistical analysis was performed using unpaired two tailed Student t-test, one sample t-test or Mann–Whitney U test using GraphPad Prism version 8. The significance of mean comparison is annotated as follows: ns, non-significant (p > 0.05), *p ≤ 0.05, **p ≤ 0.01, ***p ≤ 0.001, and ****p ≤ 0.0001.

## Acknowledgements

We thank Krzysztof Kolmus for performing the GO analysis of AXL interactors. We also thank Magdalena Banach-Orłowska, Małgorzata Maksymowicz and Marta Kaczmarek for critical reading of the manuscript. We are grateful to Raymond B. Birge for providing a plasmid encoding GAS6. We acknowledge the support of Katarzyna Mleczko-Sanecka and members of her laboratory in designing a strategy for the generation of CRISPR-Cas9 knockout cell lines and sharing control non-targeting sgRNA.

## Author contributions

DZB conceived and designed the research with support from MM, KOS and HS. DZB performed and analyzed most of the experiments with support from AP and KK (cellular and biochemical experiments), KJ (high resolution confocal microscopy and live-cell imaging) and KOS (live-cell imaging), MBO and KP (flow cytometry). AP performed the BioID and invasion spheroid assays with support from DZB. DZB wrote the manuscript with support from MM, AP, KK and KJ. All authors approved the manuscript.

## Expanded View Figure legends

**Fig EV1.**
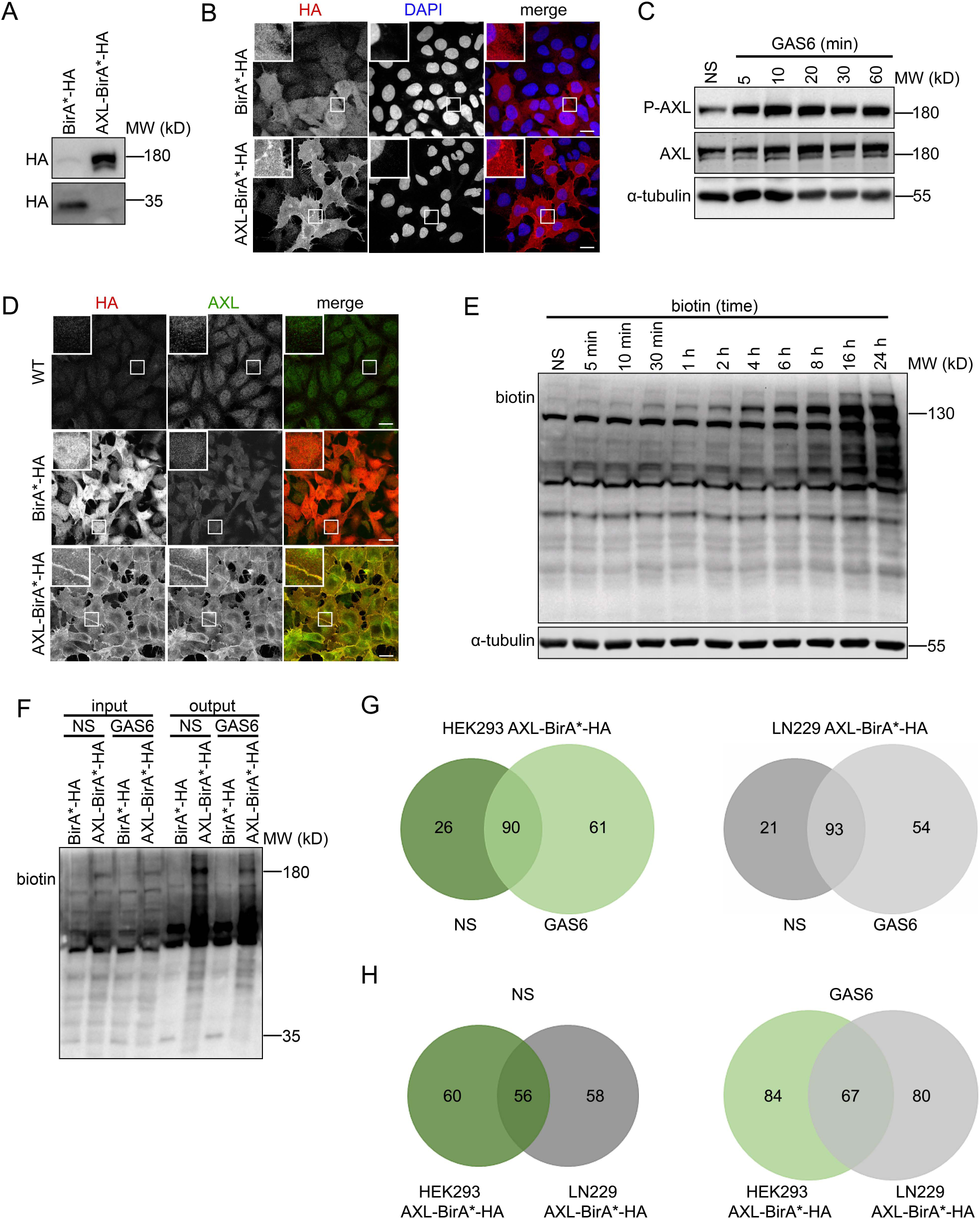
Identification of AXL interactors using the BioID assay. **A** Western blot showing expression of BirA*-HA and AXL-BirA*-HA in HEK293 cells. Antibodies recognizing HA were used. **B** Confocal images showing the localization of BirA*-HA and AXL-BirA*-HA in HEK293 cells after transient transfection with pcDNA3.1-BirA*-HA and pcDNA3.1-AXL-BirA*-HA plasmids, respectively. Cells were fixed and stained with DAPI (blue) and antibodies recognizing HA (red). **C** Western blot showing AXL-BirA*-HA phosphorylation (P-AXL, Y702). HEK293 cells expressing AXL-BirA*-HA were stimulated with GAS6 for the indicated time periods. α-tubulin was used as a loading control. **D** Confocal images showing the localization of BirA*-HA and AXL-BirA*-HA in clonally selected HEK293 cells. Cells were fixed and stained with antibodies recognizing HA (red) and AXL (green). **E** Western blot showing protein biotinylation in LN229 AXL-BirA*-HA cells incubated with biotin for the indicated time periods. Antibodies against biotin were used for blotting. α-tubulin was used as a loading control. **F** Western blot showing biotinylation status of proteins before (input) and after (output) pull-down with streptavidin-coated magnetic beads. HEK293 cells expressing BirA*-HA or AXL-BirA*-HA were incubated with biotin for 24 h, in the presence or absence of GAS6, lysed and processed as described in the Methods section. Antibodies against biotin were used for blotting. **G** Venn diagrams depicting the numbers of AXL-interacting proteins identified in HEK293 (left panel) or LN229 cells (right panel) in the absence or presence of GAS6. **H** Venn diagrams showing the comparison of numbers of AXL interactors identified in non-stimulated (left panel) or GAS6-stimulated (right panel) HEK293 and LN229 cells expressing AXL-BirA*-HA. Data information: Insets in confocal images are magnified views of boxed regions in the main images. Scale bars: 20 μm. NS-non-stimulated cells, GAS6-GAS6-stimulated cells, WT-wild type cells.

**Fig EV2.**
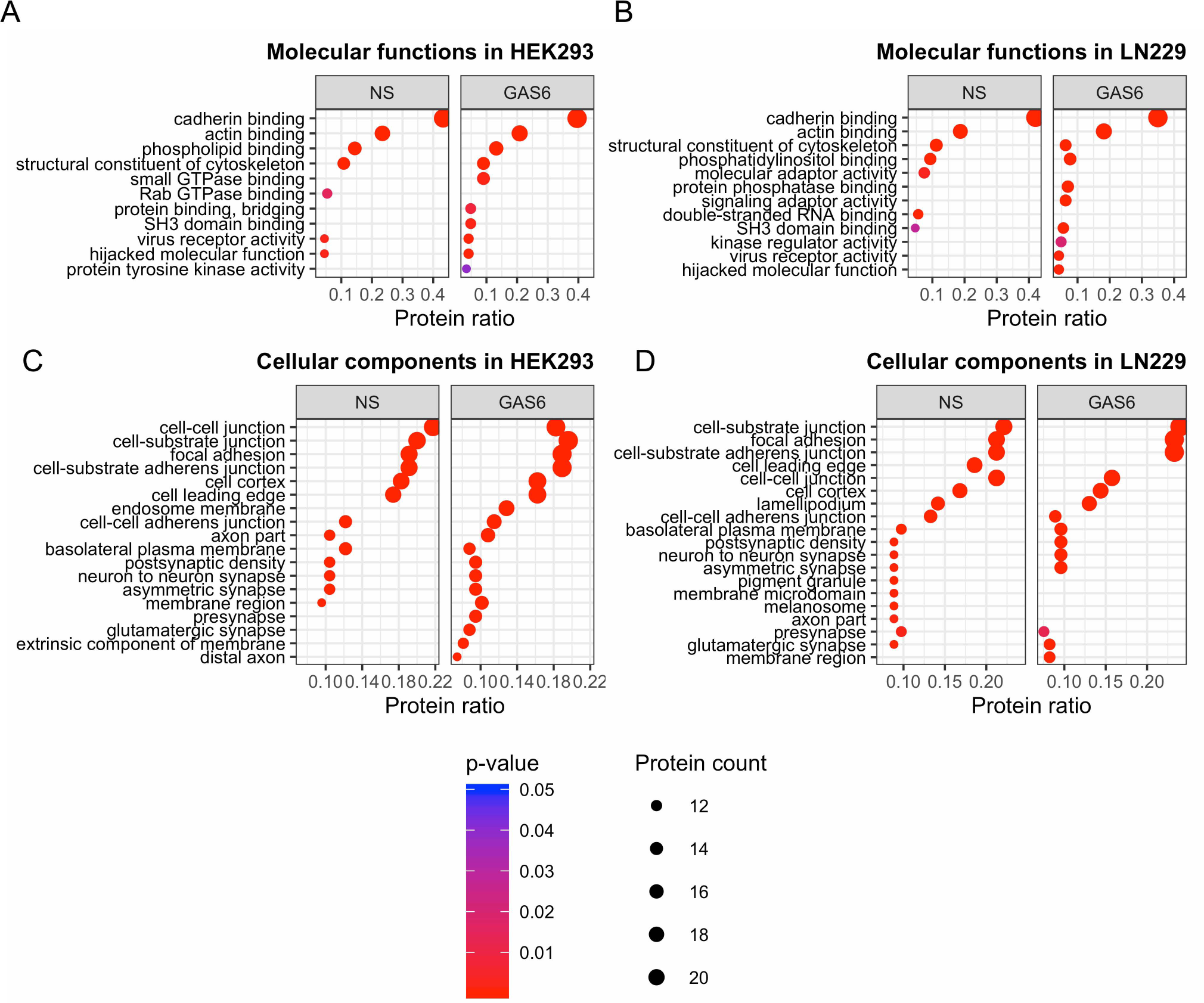
GO analyses of the BioID hits. **A, B** Graphs showing the GO analysis of molecular functions for the BioID hits identified in HEK293 (A) and LN229 cells (B). **C, D** Graphs showing the GO analysis of cellular components for the BioID hits identified in HEK293 (C) and LN229 cells (D).

**Fig EV3.**
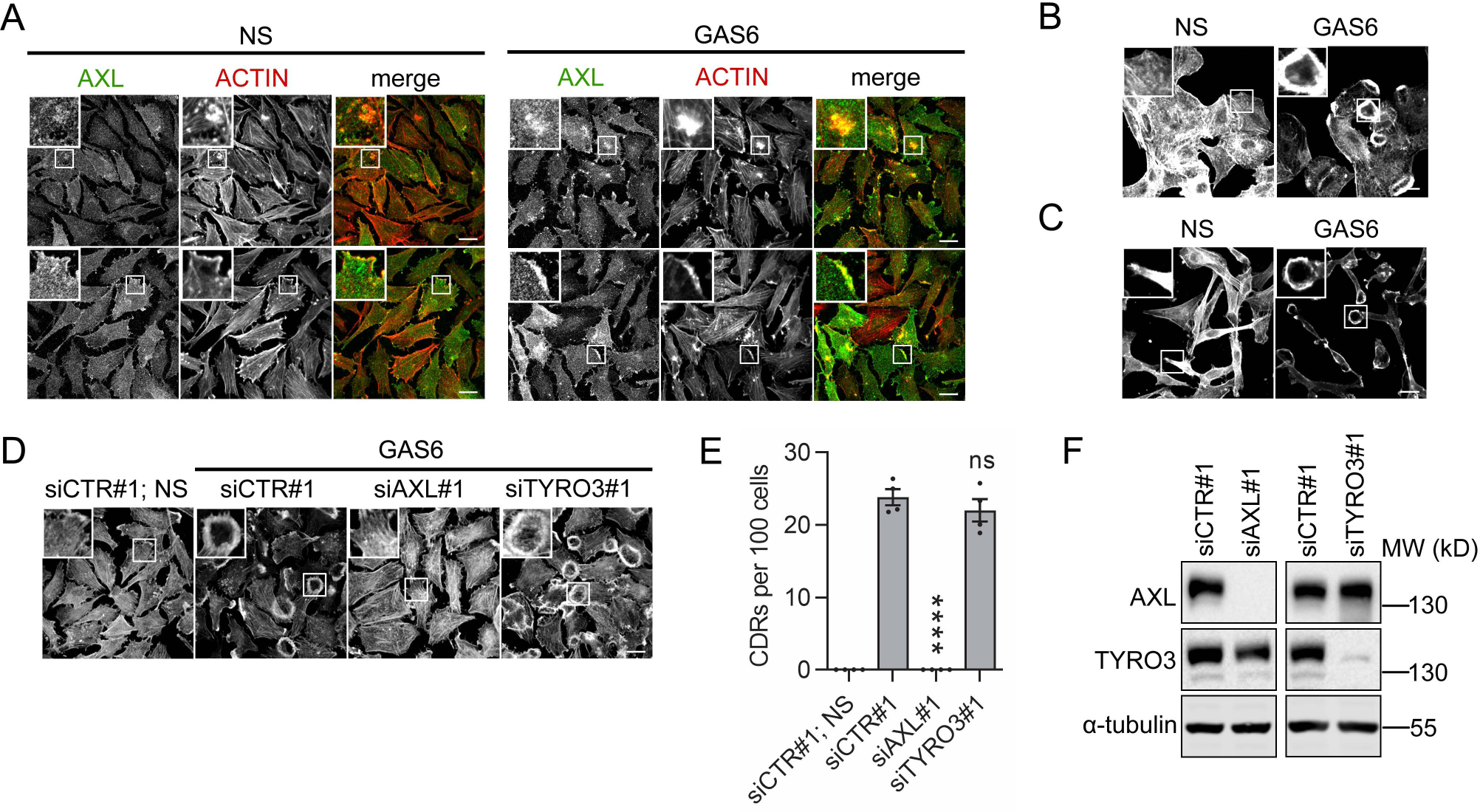
GAS6 induces the formation of CDRs which depends on AXL. **A** Confocal images showing the accumulation of AXL in cell regions, where actin is highly concentrated (top panels) including the area of lamellipodium (bottom panel) in non-stimulated and GAS6-treated LN229 cells. Serum-starved LN229 were stimulated with GAS6 for 5 min, fixed and co-stained with antibodies for AXL (green) and phalloidin to visualize actin (red). **B, C** Confocal images showing GAS6-induced CDR formation in A549 and MDA-MB-231 cells, respectively. Serum-starved cells were stimulated with GAS6 for 10 min, fixed and actin was stained with phalloidin. **D** Confocal images showing CDR formation upon siRNA-mediated depletion of AXL or TYRO3 in LN229 cells transfected with non-targeting siRNA (siCTR#1), siRNA targeting *AXL* (siAXL#1) or *TYRO3* (siTYRO3#1). 72 h after transfection serum-starved cells were stimulated with GAS6 for 10 min, fixed and actin was stained with phalloidin. **E** Quantification of data shown in (D). Approximately 150 cells were counted per experiment. Each dot represents data from one independent experiment whereas bars represent the means ± SEM, n=4. Student’s unpaired t-test, ****p≤0.0001. ns-non-significant (p>0.05). **F** Western blot showing the efficiency of siRNA-mediated AXL and TYRO3 depletion (as in D). α-tubulin was used as a loading control. Data information: Insets in confocal images are magnified views of boxed regions in the main images. Scale bars: 20 μm. NS-non-stimulated cells, GAS6-GAS6-stimulated cells.

**Fig EV4.**
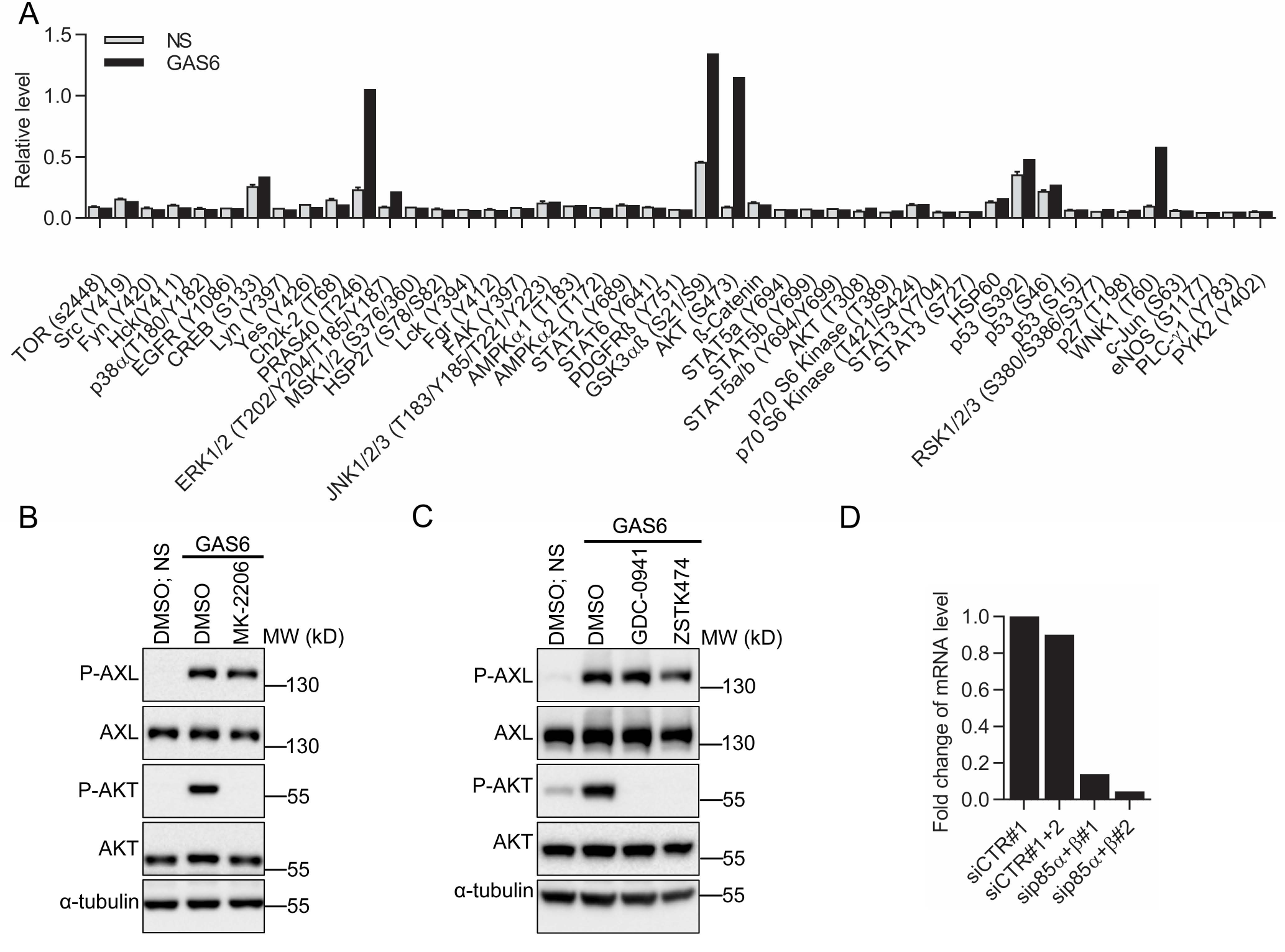
PI3K mediates GAS6-induced CDR formation. **A** The densitometric analysis of the phospho*-*kinase array data shown in Fig 3A. Phosphorylation of 43 human kinases and total amounts of 2 related proteins (all listed on the x-axis) were tested. The levels of phosphorylated forms of the indicated proteins were normalized to appropriate positive controls. **B** Western blot showing GAS6-induced AXL (P-AXL, Y702) and AKT (P-AKT, S473) phosphorylation upon AKT inhibitor. Serum-starved LN229 cells were pretreated with MK-2206 for 30 min prior to stimulation with GAS6 for 10 min. α-tubulin was used as a loading control. **C** Western blot showing GAS6-induced AXL (P-AXL, Y702) and AKT (P-AKT, S473) phosphorylation upon treatment with PI3K inhibitors. Serum-starved LN229 cells were pretreated with GDC-0941 or ZSTK474 for 30 min prior to stimulation with GAS6 for 10 min. α-tubulin was used as a loading control. **D** Graph showing the efficiency of siRNA-mediated silencing of genes encoding p85α and p85β. LN229 cells were transfected with non-targeting siRNA (siCTR#1), the combination of non-targeting siRNAs (siCTR#1+siCTR#2), two combinations of siRNAs targeting p85α and p85β (sip85α#1+p85β#1 and sip85α#2+p85β#2). 72 h after transfection expression level of genes encoding p85α and p85β was measured by qRT-PCR. Values are presented as fold change with respect to siCTR#1.

**Fig EV5.**
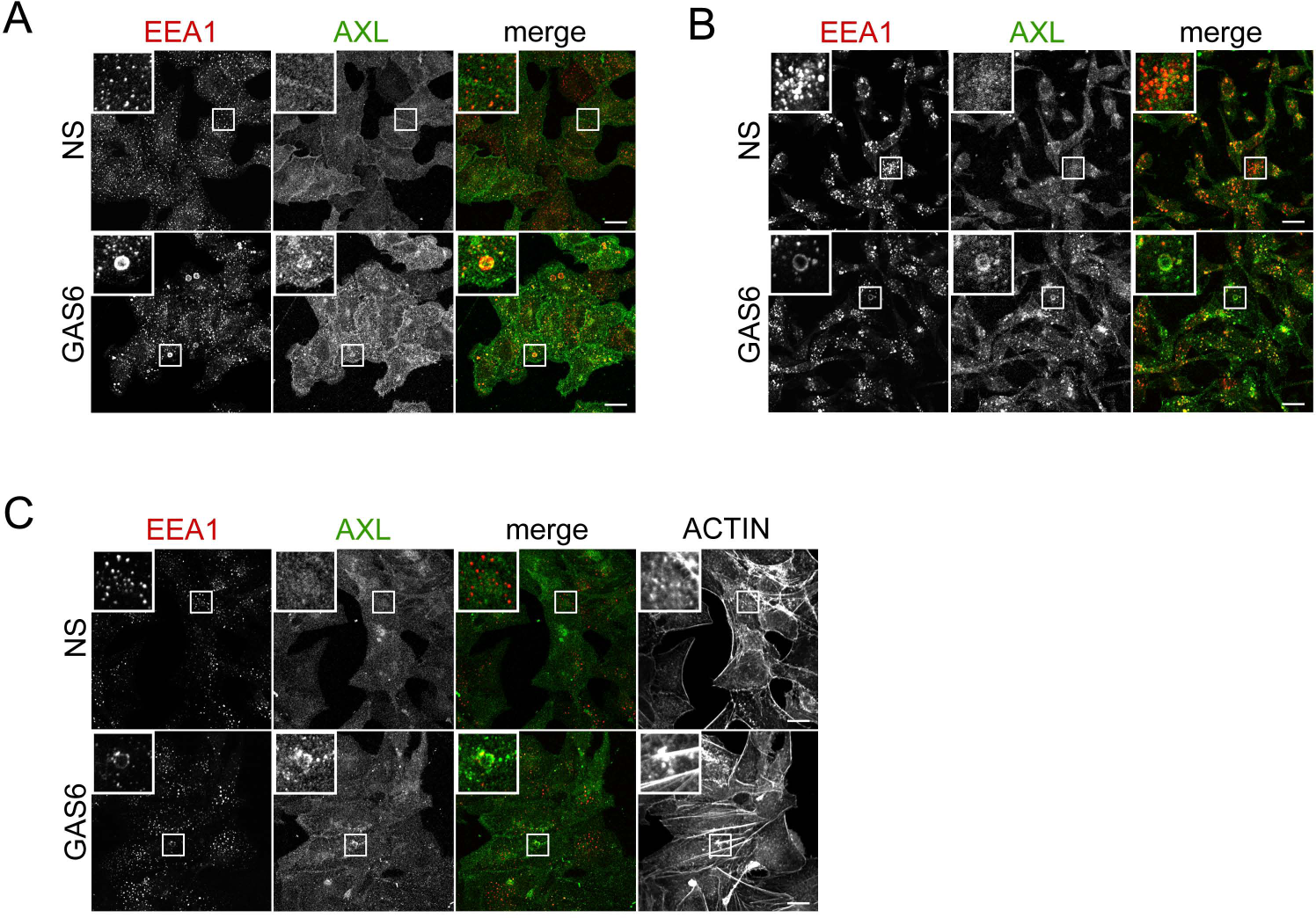
GAS6 stimulates macropinosome formation in different cell lines. **A-C** Confocal images showing GAS6-induced macropinosome formation in A549 (A), MDA-MB-231 (B) and SKOV3 (C) cells, respectively. Serum-starved cells were stimulated with GAS6 for 10 min, fixed and co-stained with antibodies for AXL (green) and EEA1 (red). Additionally actin was stained with phalloidin (C). Data information: Insets are magnified views of boxed regions in the main images. Scale bars: 20 μm. NS-non-stimulated cells, GAS6-GAS6-stimulated cells.

**Fig EV6.**
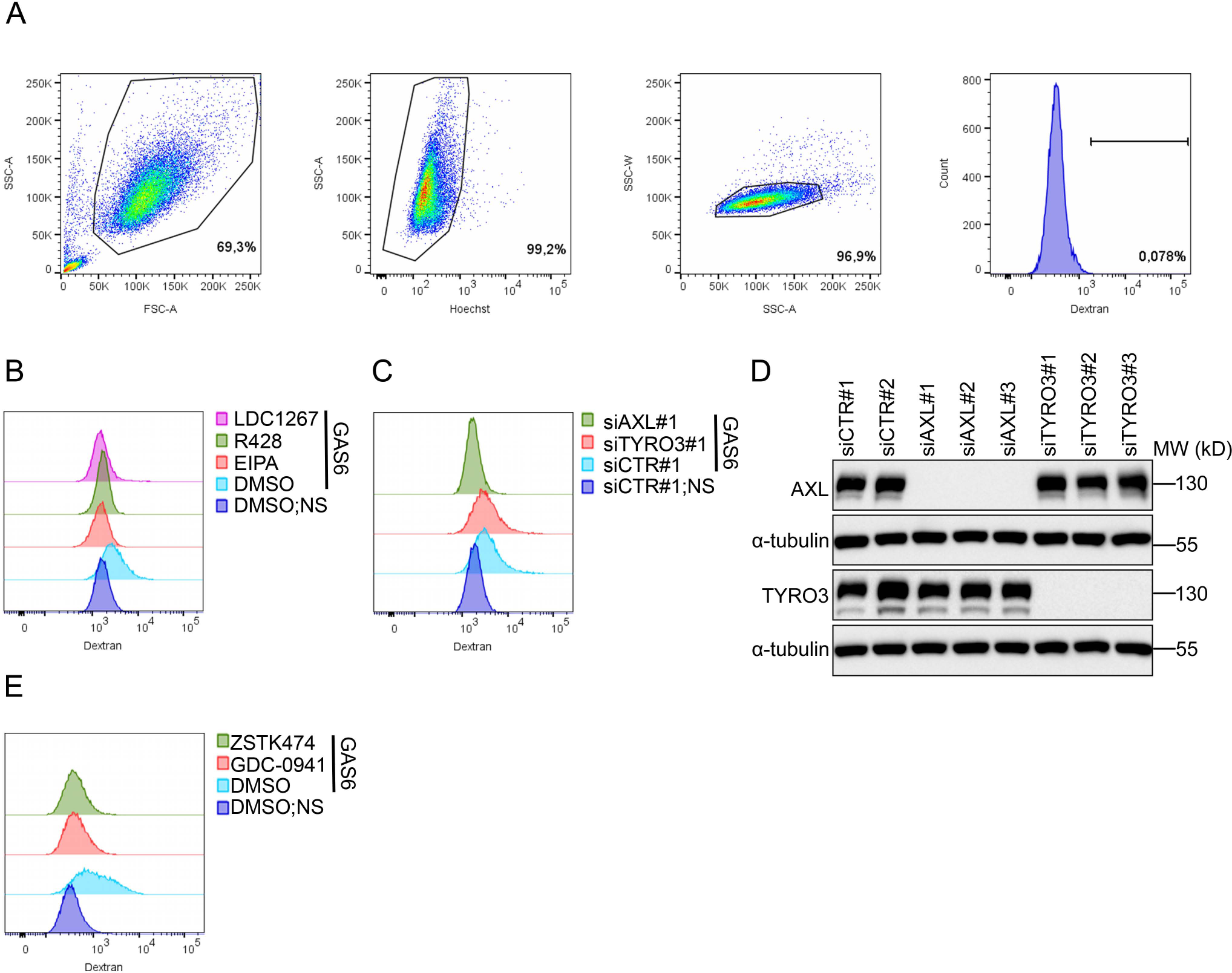
GAS6-induced macropinocytic internalization of dextran is blocked by EIPA, AXL and PI3K inhibitors. **A** Examples of gating strategy for flow cytometric analysis shown in Fig 5C-E, including cell gate (FSC vs SSC), live cell discrimination gate (Hoechst vs SSC), singlet discrimination gate (SSC-A vs SSC-W), and cells that internalized fluorescent dextran. **B** Overlay histograms showing fluorescence signal of dextran internalized by serum-starved LN229 cells pretreated with macropinocytosis (EIPA) and AXL inhibitors (R428 and LDC1267) prior to GAS6 stimulation. The histograms illustrate an example experiment included in the analyses shown in Fig 5C. **C** Overlay histograms showing fluorescence signal of dextran internalized by GAS6-stimulated LN229 cells upon siRNA-mediated depletion of AXL or TYRO3. The histograms illustrate an example experiment included in the analyses shown in Fig 5D. **D** Western blot showing the efficiency of siRNA-mediated AXL and TYRO3 depletion (as in Fig 5D). α-tubulin was used as a loading control. **E** Overlay histograms showing fluorescence signal of dextran internalized by LN229 cells pretreated with and PI3K inhibitors prior to GAS6 stimulation. The histograms illustrate an example experiment included in the analyses shown in Fig 5E.

**Fig EV7.**
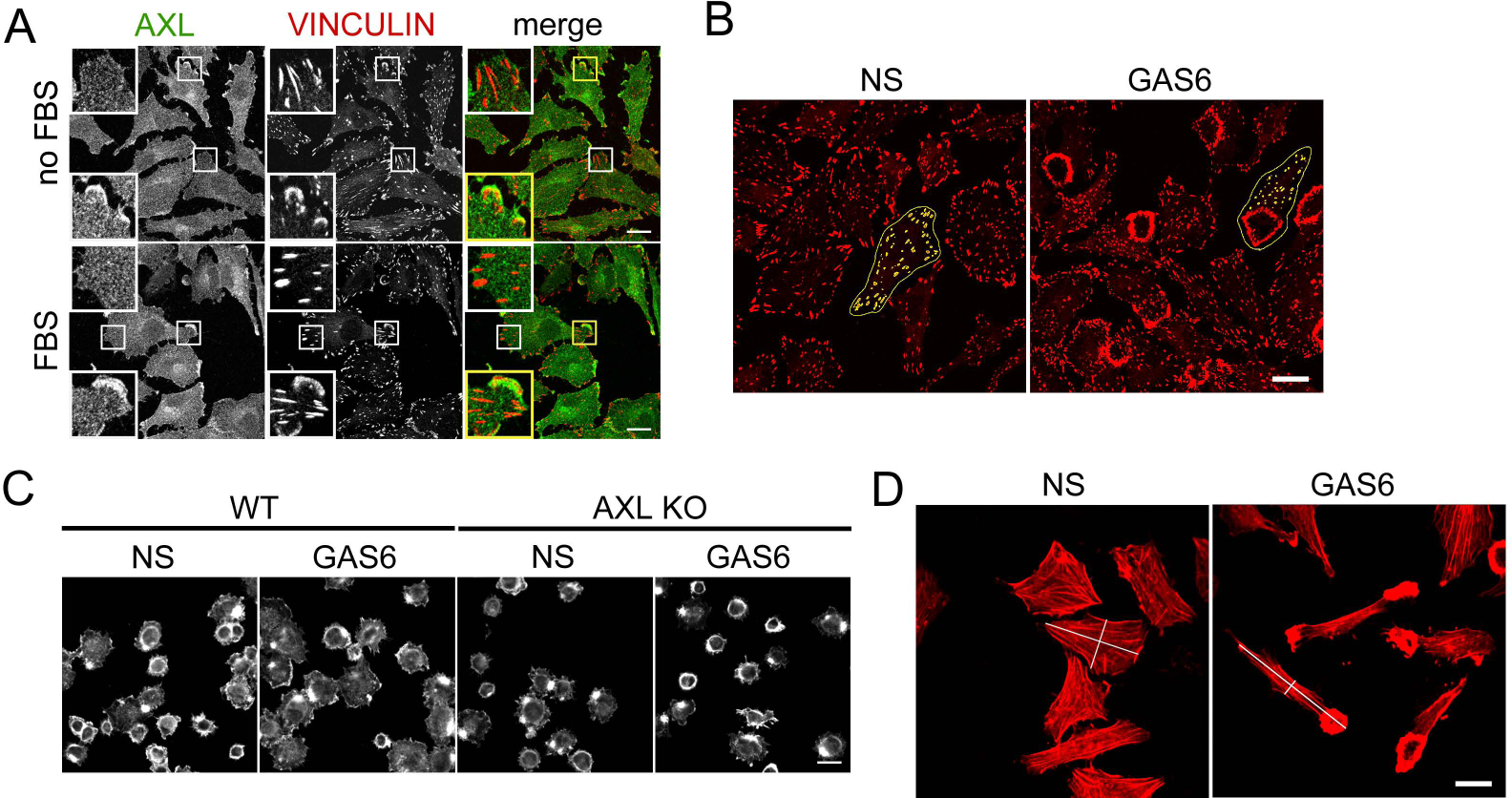
AXL accumulates on lamellipodia but not FAs (A). GAS6-induced CDRs reduce the number of FAs and GAS6 increases cell spreading and cell elongation index (B-D). **A** Confocal images showing the accumulation of AXL and vinculin on lamellipodia (yellow inset) and lack of AXL accumulation on FAs (white inset) in LN229 cells grown without serum (no FBS) or in full medium (FBS). Cells were fixed and stained with antibodies for AXL (green) and vinculin (red). **B** Confocal images illustrating examples of cells (contoured) analyzed in Fig 6F and G for number and average area of FAs (marked in yellow) in non-stimulated and GAS6-treated LN229 cells forming CDRs. **C** Example confocal images that were analyzed in Fig 6H to measure spreading of wild type (WT) and AXL KO (gAXL#2) LN229 cells following GAS6 stimulation. **D** Example confocal images used in Fig 6I to calculate the cell elongation index of non-stimulated and GAS6-stimulated LN229 cells with CDRs. Examples of the major and the minor axes are drawn. Data information: Insets are magnified views of boxed regions in the main images. Scale bars: 20 μm. NS-non-stimulated cells, GAS6-GAS6-stimulated cells, WT-wild type.

**Fig EV8.**
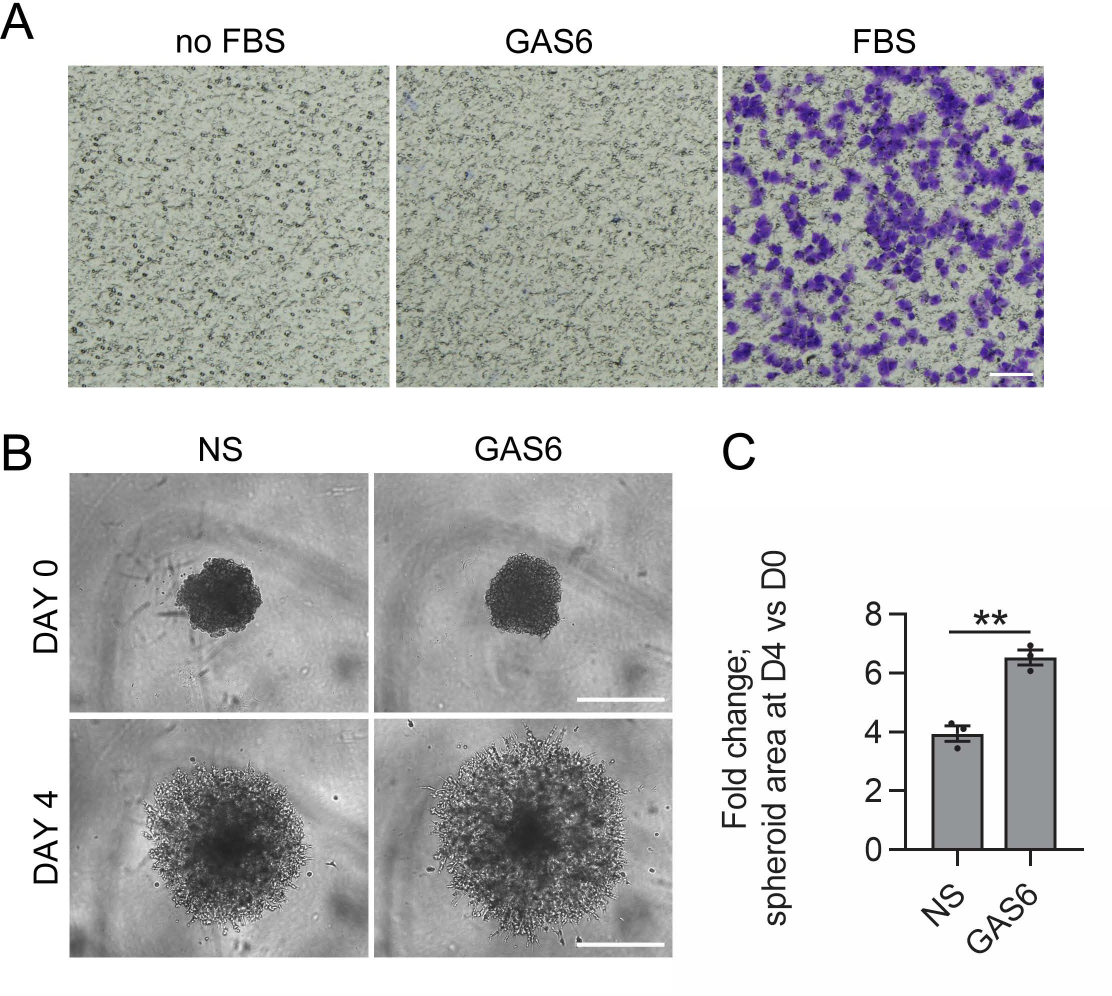
GAS6 is not a chemoattractant for LN229 cells (A). GAS6 stimulates invasion of MDA-MB-231 spheroids (B-C). **A** Images showing LN229 cells that migrated through microporous membrane in a Transwell migration assay. Cells were seeded in serum-free medium in the upper chamber, whereas serum-free medium (no FBS, negative control), serum-free medium with GAS6 (GAS6) or medium with 10% FBS (FBS, positive control) were added to the lower chamber. Migrated cells were fixed and stained with crystal violet. Representative images from two independent experiments are shown. Scale bars: 100 μm. **B** Images showing GAS6-stimulated invasion of MDA-MB-231 spheroids into Matrigel. Spheroids were grown in the presence or absence of GAS6 in Matrigel for 4 days. Scale bars: 500 μm. **C** Quantification of data shown in (B). The area of spheroids was measured by ImageJ software. Data are expressed as fold changes of the spheroid area on the 4^th^ day (DAY 4, D4) with respect to the spheroid area before Matrigel addition (DAY 0, D0). Each dot represents data from one independent experiment whereas bars represent the means ± SEM, n=3. Student’s unpaired t test, **p<0.01. NS-non-stimulated cells, GAS6-GAS6-stimulated cells.

Movie EV1 and 2

**GAS6 induces membrane ruffling and macropinosome formation.** Serum-starved LN229 cells expressing the plasma membrane tethered mCherry (MyrPalm-mCherry) were monitored by spinning disk confocal microscopy for 5 min before and 30 min after addition of GAS6. The images were acquired every 30 s. Scale bars: 20 μm.

Movie EV3 and 4

**GAS6 induces CDRs and macropinosomes, the latter formed upon CDR closure.** Serum-starved LN229 cells expressing MyrPalm-mCherry and LifeAct-mNeonGreen were monitored by fluorescence microscopy for 5 min before and 20 min after addition of GAS6. The images were acquired every 30 s. Scale bars: 10 μm.

## EV Data legends

**Data EV1.** The lists of AXL interactors identified in non-stimulated (NS) and GAS6-treated (GAS6) HEK293 cells. Proteins fulfilling the following criteria were considered as AXL proximity interactors: they were identified in at least 2 out of 3 experiments, with ≥ 2 peptides (Number of significant sequences) at least in one experiment, and had three times higher Mascot score (Score) in AXL-BirA*-HA samples in comparison to the control BirA*-HA samples (Ratio of scores). If no peptides were identified for a given hit in control samples, the ratio of scores was marked as not applicable (n/a). Data obtained in three independent experiments (I, II, III) are shown.

**Data EV2.** The lists of AXL interactors identified in non-stimulated (NS) and GAS6-treated (GAS6) LN229 cells. Proteins fulfilling the following criteria were considered as AXL proximity interactors: they were identified in at least 2 out of 3 experiments, with ≥ 2 peptides (Number of significant sequences) at least in one experiment, and had three times higher Mascot score (Score) in AXL-BirA*-HA samples in comparison to the control BirA*-HA samples (Ratio of scores). If no peptides were identified for a given hit in control samples, the ratio of scores was marked as not applicable (n/a). Data obtained in three independent experiments (I, II, III) are shown.

## EV Tables and their legends

Table EV1. AXL proximity interactors (detected in HEK293 and/or LN229 cells) previously reported to localize and/or be involved in the formation of CDRs.

Table EV1 is attached as separate file.

**Table EV2.**
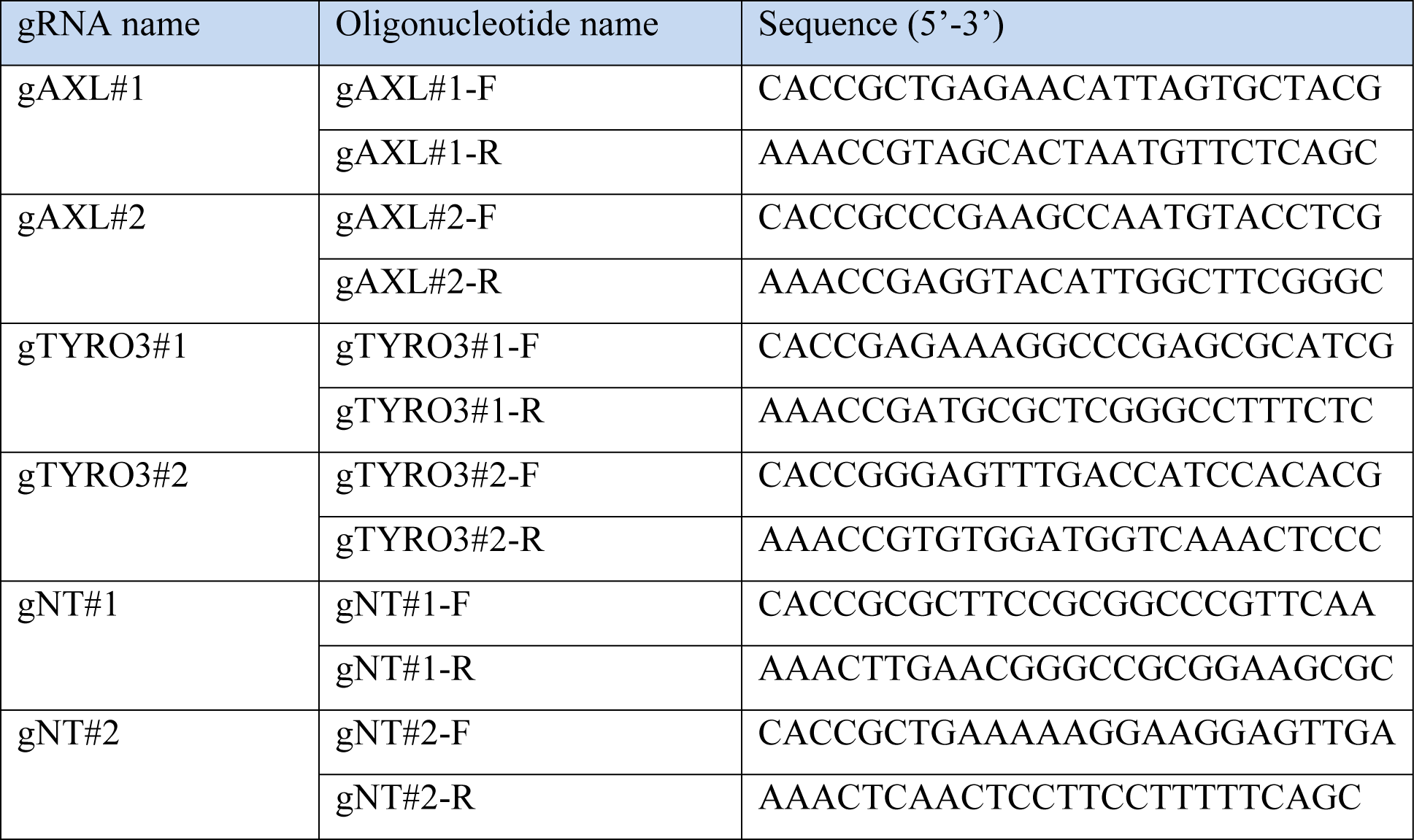
gRNAs used for CRISPR-Cas9-mediated gene inactivation. For each gene, two 20-bp-long single-guide RNA (gRNAs) were selected from the Brunello library (Doench et al, 2016), and appropriate pairs of DNA oligonucleotides were designed as described elsewhere (Sanjana et al, 2014).

**Table EV3.**
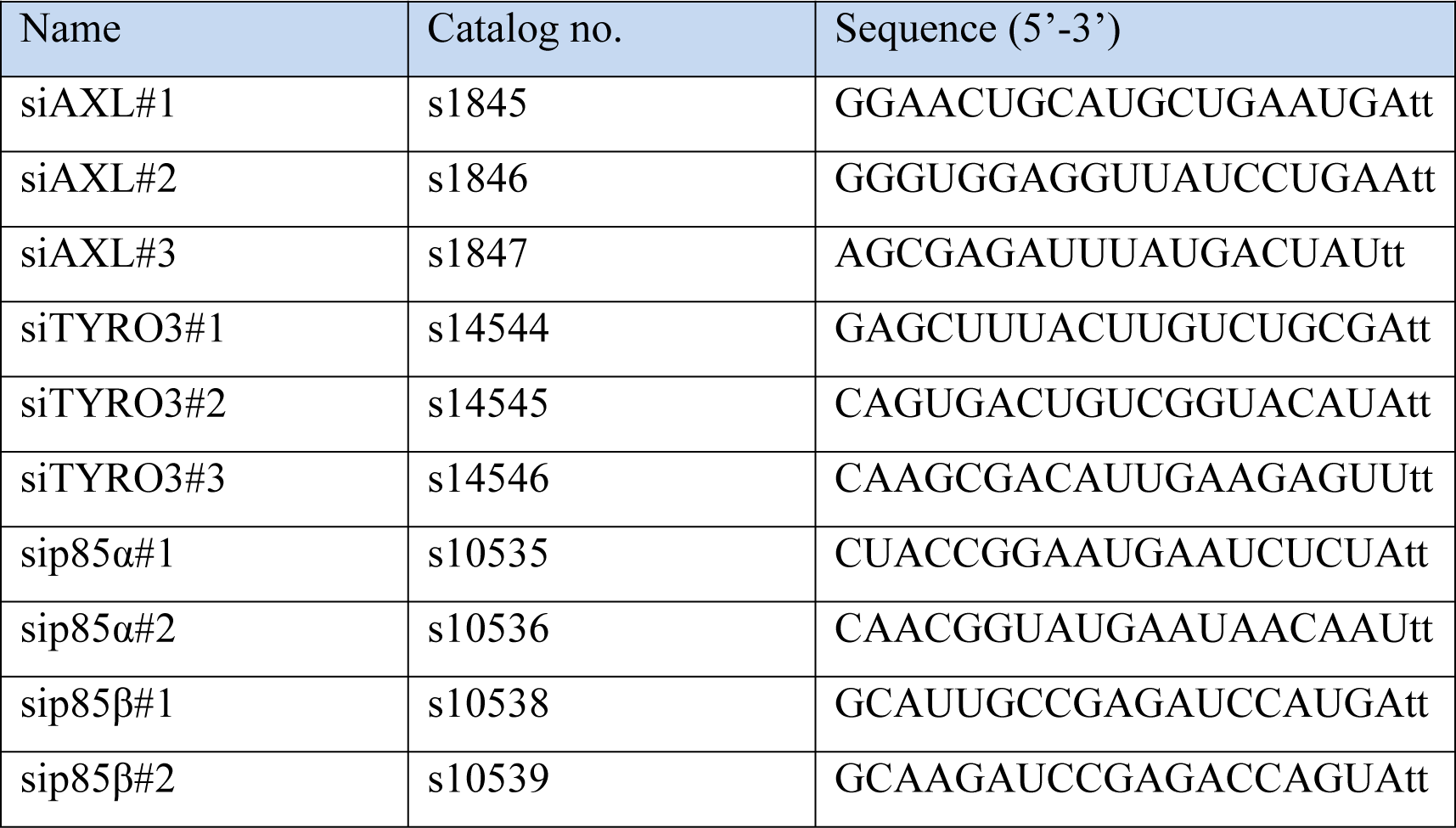
Small interfering RNA (siRNA) oligonucleotides. Ambion Silencer Select siRNAs were from Thermo Fisher Scientific. As negative control, non-specific Silencer Select siRNA oligonucleotides (siCTR#1, 4390846 and siCTR#2, AM4615) were used.

**Table EV4.**
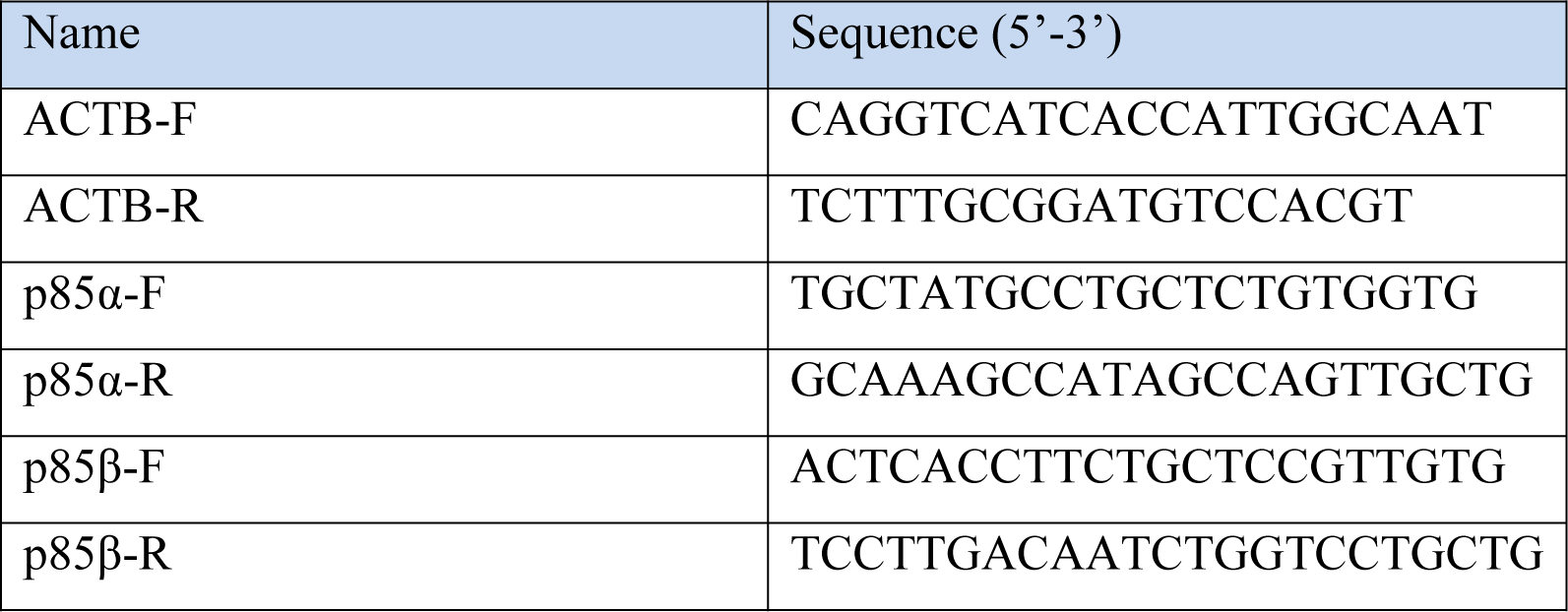
Primers used for qRT-PCR. Primers used for qRT-PCR were designed using QuantPrime and custom-synthesized by Sigma-Aldrich.

